# Induction of interhemispheric visual plasticity by paired associative stimulation

**DOI:** 10.64898/2026.10.05.756694

**Authors:** Francesca Crespi, Giacomo Guidali, Eleonora Arrigoni, Nadia Bolognini

**Affiliations:** PhD program in Neuroscience, School of Medicine and Surgery, University of Milano-Bicocca, Monza, IT; Department of Psychology, University of Milano-Bicocca, Milano, IT; Laboratory of Neuropsychology, IRCCS Istituto Auxologico Italiano, Milano, IT

**Keywords:** primary visual cortex, associative plasticity, transcranial magnetic stimulation, visual processing, interhemispheric transfer

## Abstract

Unveiling the plastic mechanisms of the primary visual cortex (V1) is crucial for understanding how visual processing can change and adapt, ultimately informing interventions to promote functional recovery following visual system damage. Paired associative stimulation (PAS) protocols can induce associative plasticity within the human V1. However, previous paradigms used full-field visual stimulation, engaging both hemispheres simultaneously, leaving unclear whether plasticity can be selectively targeted to a single visual pathway and whether it can modulate the non-stimulated hemisphere.

Across three experiments in healthy participants, we investigated the neurophysiological, behavioural, and interhemispheric outcomes of a novel lateralized visual PAS protocol (vPAS). Pattern-reversal checkerboards were presented in a single visual hemifield and time-locked to transcranial magnetic stimulation (TMS) pulses delivered over the contralateral V1. *Experiment 1* assessed neurophysiological effects by recording hemifield-lateralized visual-evoked potentials (VEPs) and testing three inter-stimulus intervals (ISIs: 75, 100, and 125 ms) between vPAS-paired stimuli. *Experiment 2* evaluated perceptual changes in contrast detection, while *Experiment 3* examined vPAS hemispheric specificity and aftereffects on interhemispheric visuomotor transfer.

Only the 75-ms ISI consistently modulated visual processing in the non-conditioned hemisphere, reducing N75 amplitude and improving contrast detection for visual stimuli presented contralateral to the TMS side. Longer ISIs were not effective. Furthermore, this protocol altered interhemispheric transfer exclusively when the left, but not the right, V1 was targeted during vPAS.

In conclusion, lateralized vPAS induces timing-dependent changes beyond the stimulated cortex, suggesting that associative plasticity can engage contralateral visual circuits and modulate visual processing.

## 1. INTRODUCTION

Uncovering the plastic properties of the human primary visual cortex (V1) is crucial for understanding how visual processing changes and adapts, ultimately informing interventions to promote functional recovery following visual system damage (e.g., Hooks & Chen, 2007; Spolidoro et al., 2009; Wandell & Smirnakis, 2009). Non-invasive brain stimulation techniques, such as transcranial magnetic stimulation (TMS), have emerged as powerful tools for probing and shaping V1 plasticity. TMS, in particular, can effectively modulate visual cortical excitability, influencing perceptual functions such as contrast sensitivity, motion detection, and phosphene perception, as well as visual-evoked potentials (VEPs) (Antal et al., 2006; Bello et al., 2023; Bocci et al., 2014; Siebner et al., 2022).

Among TMS protocols, paired associative stimulation (PAS) has been shown to effectively induce associative plasticity across various cortical networks, including the visual system (Guidali et al., 2021b, 2021a; Suppa et al., 2017). PAS relies on the repeated, time-locked pairing of a peripheral stimulation and a TMS pulse delivered over the corresponding cortical area. The temporal interval between the two inputs is a key factor for PAS effectiveness: when the sensory signal arrives at the cortex immediately before the TMS pulse, achieving temporal contingency between endogenous and exogenous activation, long-term potentiation (LTP)-like effects are typically induced; conversely, delivering the TMS pulse long before sensory arrival can result in long-term depression (LTD)-like effects (Arrigoni et al., 2026; Stefan et al., 2000; Wolters et al., 2003). Over the past decade, PAS protocols pairing visual stimuli with TMS has been successfully applied in healthy and clinical populations to investigate cross-modal and visuomotor interactions (e.g., Guidali et al., 2020, 2023; Guidali & Bolognini, 2025; Maddaluno et al., 2020; Picardi et al., 2025; Suppa, Li Voti, et al., 2015; Suppa, Rocchi, et al., 2015; Wolfe et al., 2021; Zazio et al., 2019, 2026). Similarly, cortico-cortical PAS – pairing TMS pulses over two brain areas of the visual stream – has been used to modulate low- and high-order visual processing (e.g., Bevilacqua et al., 2023, 2025; Borgomaneri et al., 2023; Chiappini et al., 2018, 2022; Romei et al., 2016; Tarasi et al., 2024).

Regarding visual protocols specifically targeting V1 (i.e., vPAS), the standard approach consists of pairing full-field pattern-reversal visual stimulation with TMS pulses delivered over V1, time-locked to the latency of early VEP components. This protocol can induce timing-dependent changes in VEP amplitude and habituation, providing evidence that PAS can induce associative plasticity in V1 in both healthy individuals (Ranieri et al., 2019) and clinical populations (Abagnale et al., 2021). However, its reliance on full-field visual stimulation engages visual cortices bilaterally. Consequently, it remains unknown whether associative plasticity can be induced selectively in one hemisphere. Resolving this question is essential for characterizing the local effects of vPAS and determining whether and how they extend beyond the stimulated visual cortex, ultimately supporting the development of this protocol for targeted clinical applications in patients with unilateral visual field defects (e.g., hemianopia).

Building on these premises, the present study investigated the neurophysiological, perceptual, and interhemispheric impact of a novel, lateralized vPAS protocol pairing pattern-reversal black-and-white checkerboards presented within a single visual hemifield with TMS pulses delivered over the contralateral V1. Across three experiments, we systematically evaluated: (i) neurophysiological changes in hemifield-lateralized VEPs (*Experiment 1*), (ii) perceptual modifications in visual contrast sensitivity (*Experiment 2*), and (iii) hemispheric specificity alongside inter-hemispheric visuomotor transfer dynamics (*Experiment 3*).

In particular, in *Experiment 1*, pattern-reversal VEPs were recorded from both visual fields before and after three vPAS protocols differing in their interstimulus interval (ISI). Identifying the optimal ISI is indeed a fundamental requirement for establishing PAS efficacy (Guidali et al., 2021b; Suppa et al., 2017). We selected three ISIs – 75, 100, and 125 ms – based on the mean latencies of the main pattern-reversal VEP component (Blumenhardt & Halliday, 1979; Kurita-Tashima et al., 1991; Shigeto et al., 1998). We chose the 75-ms ISI to align with the N75 component, which reflects the early cortical response primarily generated within the striate cortex. The 100-ms ISI targeted the P100 component, generated by combined striate and extra-striate activity, while the 125-ms ISI targeted later processing stages with higher extra-striate involvement (Boehler et al., 2008; Di Russo et al., 2001; Pratt, 2011; Šuštar Habjan et al., 2025). We hypothesized that vPAS-75 and vPAS-100 would fall within the optimal temporal window needed for the visual input to interact contingently with the TMS pulse in V1, inducing LTP-like modulation of VEP amplitudes. Conversely, the 125-ms ISI was expected to be less favourable for plasticity induction, providing a control condition to assess the protocol’s timing dependence.

In *Experiment 2*, informed by the electrophysiological findings of *Experiment 1*, we assessed whether vPAS modulates visual perception, specifically visual contrast sensitivity. Because V1 neurons play a central role in encoding stimulus contrast and supporting contrast-dependent orientation judgments (Albrecht & Hamilton, 1982; Boynton et al., 1999; Himmelberg et al., 2022), we hypothesized that lateralized vPAS would induce hemifield-specific enhancements in contrast detection.

Finally, in *Experiment 3*, we examined whether vPAS effects exhibit hemispheric asymmetry and whether local plastic induction propagates to broader interhemispheric visuomotor networks. Converging behavioural, electrophysiological, neuroimaging, and TMS evidence indicates that the left and right visual areas exhibit distinct functional dynamics, including symmetric interhemispheric transfer, complementary specialization for local *versus* global visual processing, and distinct spatiotemporal signal propagation following occipital stimulation (Brown et al., 1994; Hellige, 1996; Marzi, 1999; Siviero et al., 2023). While *Experiments 1* and *2* exclusively targeted the left V1, *Experiment 3* directly tested hemispheric specificity by engaging the right V1 (reversing both the presentation hemifield and TMS site). Additionally, to test whether induced plasticity modulates interhemispheric interactions, participants underwent the Poffenberger paradigm (Marzi, 1999; Poffenberger, 1912), in which reaction times to lateralized visual stimuli provide a well-established measure of interhemispheric transfer efficiency (e.g., Braun, 1992; Chaumillon et al., 2014; Wolski & Asanowicz, 2023).

## 2. EXPERIMENT 1 – Optimal timing of vPAS: neurophysiological evidence

### 2.1. Participants

Thirty-one healthy volunteers took part in *Experiment 1*. Four participants were excluded due to recording issues, and two met outlier criteria for at least one of our dependent variables (see **Statistical analysis**), leaving a final analysed sample of 25 participants (7 males, mean age ± standard deviation – SD: 21.8±3.3 years, education: 13.6±1.8 years). All participants were right-handed, according to the Edinburgh Handedness Inventory (Oldfield, 1971), and had no contraindications to TMS (Rossi et al., 2009). None had a history of neurological or ophthalmological disease, and all had normal or corrected-to-normal visual acuity. The recommended sample size was estimated *a priori* using the software G*Power (3.1.9.7; Faul et al., 2009) for a 3 (ISI: 75, 100, 125 ms) x 2 (Time: pre, post) x 2 (VF: right, left) within-group repeated-measures Analysis of Variance (rmANOVA), as our experimental design. We considered the following parameters: beta=0.95; alpha=0.05; effect size *F*=0.42 (i.e., the one found in Ranieri et al., 2019). The calculation returned a recommended sample size of n=25.

Before participating in the study, participants provided written informed consent. The protocol was conducted in accordance with the ethical standards of the Declaration of Helsinki and was approved by the Ethical Committee of the University of Milano-Bicocca (n° 750-2023).

### 2.2. Experimental procedure

Each participant underwent three experimental sessions according to the vPAS administered (i.e., vPAS-75, vPAS-100, vPAS-125). Sessions were separated by at least 48 hours and scheduled at the same time of day (i.e., morning or afternoon) to account for potential spurious effects due to circadian rhythms (Sale et al., 2007). The order of the sessions was counterbalanced across participants. The experimental procedure was the same across all sessions and consisted of the following steps: EEG cap preparation, pre-vPAS VEP recording, determination of the resting motor threshold (rMT), localization of the left V1, administration of the vPAS protocol, and post-vPAS VEP recording. On average, each session lasted approximately 2 hours. During the entire experiment, participants were seated in front of a PC monitor (diagonal: 22.3’’; aspect ratio: 16:9; refresh rate: 60 Hz) at a viewing distance of 100 cm, in an acoustically isolated room with dimmed light and a uniform luminance field of ∼5 cd/m^2^. During VEP recording and vPAS administration, participants placed their heads on a chin rest to minimize movement.

### 2.3. Visual paired associative stimulation (vPAS)

#### 2.3.1 vPAS parameters

The vPAS used in the present study was a modified version of the protocol introduced by Ranieri et al. (2019), where we paired a visual stimulus presented in the right visual hemifield with a TMS pulse delivered over the contralateral V1. In particular, TMS was delivered over left V1 at 120% of the participant’s rMT (as in Abagnale et al., 2021; Ranieri et al., 2019) following the onset of a contrast-reversing checkerboard presented in the right visual hemifield (Michelson digital contrast: 100%, mean luminance: 200 cd/m^2^, dimension: 7.44° x 7.44° visual angle, centre at 8.0° to the right of fixation). The ISI between checkerboard reversal onset and the TMS pulse varied across sessions: 75 ms (vPAS-75), 100 ms (vPAS-100), or 125 ms (vPAS-125). Each vPAS protocol comprised 180 paired stimulations delivered at 0.2 Hz; the overall protocol duration was 15 minutes (**Figure 1a**). Trial randomization, stimulus timing, and response recording were under computer control (E-Prime 3.0, Psychology Software Tool, Inc.).

**Figure 1.**
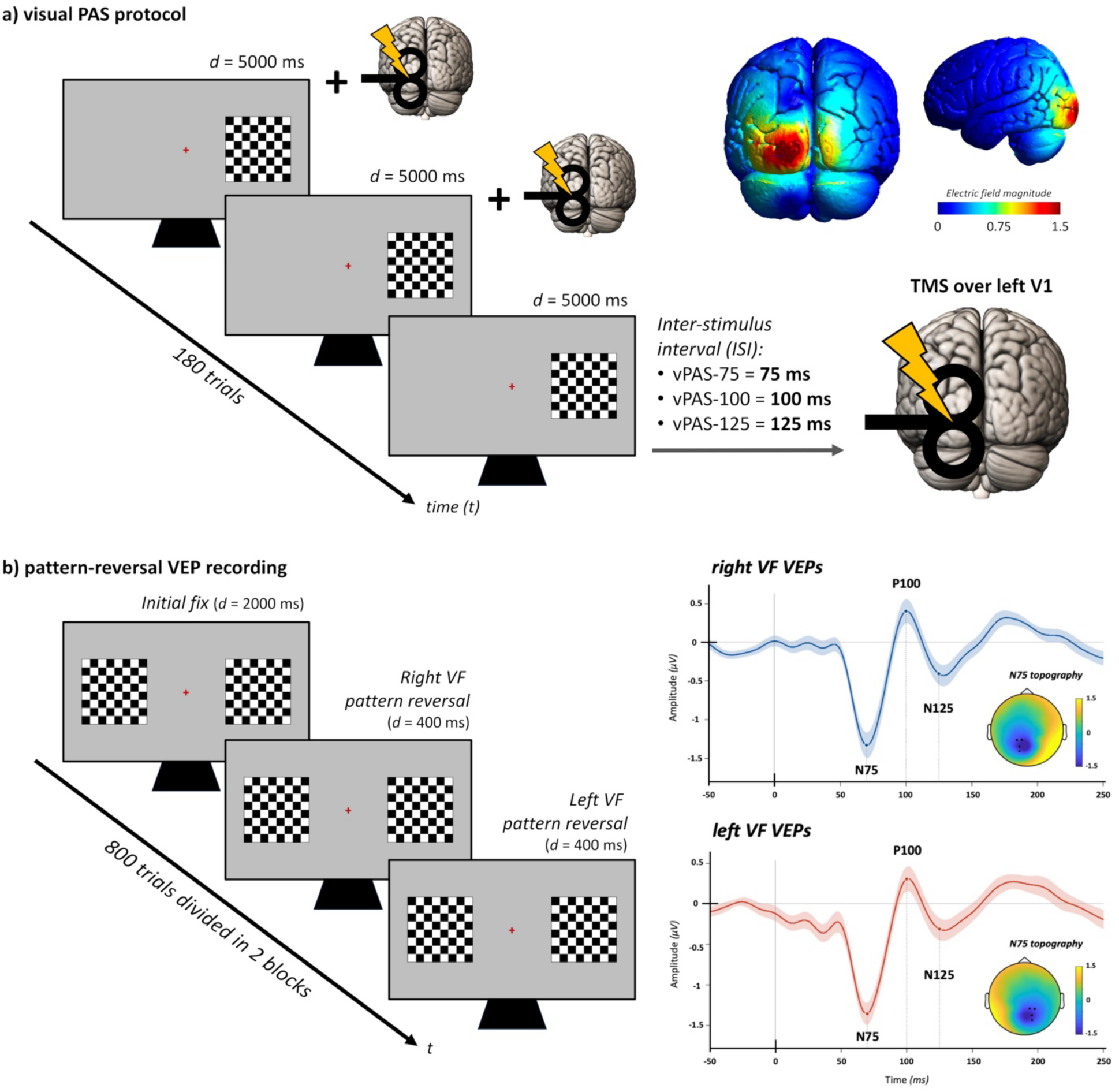
vPAS protocol (a) and pattern-reversal VEP recording (b). For (**a**), the representative electric field simulation of left V1 stimulation during vPAS administration is depicted. Simulations were performed on the standard MNI head model for TMS delivered over the O1 scalp location using a standard figure-of-eight 60-mm TMS coil model. Electric field modelling was carried out with SimNIBS (v. 4.5) (Saturnino et al., 2019). In (**b**), prototypical VEPs recorded from right and left VF from the occipital cluster in the contralateral hemisphere are depicted alongside topographies of their main components.

During vPAS, participants were instructed to maintain fixation on a red cross displayed at the centre of the screen while each checkerboard remained visible for 5 s before reversing. To ensure central fixation and maintain attention during vPAS, participants performed a detection task requiring them to respond to rare target events (18 trials) in which the fixation cross briefly changed colour from red to green for 500 ms. Participants pressed a mouse button with their right index finger upon detecting the target. Performance on this attentional task exceeded 90% accuracy across all sessions, confirming high attentional engagement during vPAS.

#### 2.3.2. Transcranial Magnetic Stimulation (TMS)

TMS was delivered using a figure-of-eight coil (diameter=60 mm) connected to a biphasic Eximia stimulator (Nexstim™, Helsinki, Finland). At the start of each vPAS session, we assessed participants’ rMT using the parameter estimation by sequential testing (PEST) method (i.e., a maximum-likelihood threshold-hunting procedure; Dissanayaka et al., 2018). This value was used to set the stimulation intensity during vPAS at 120% of rMT. We identified the motor hotspot for the right-hand first dorsal interosseous (FDI) muscle by moving the coil in 0.5-cm steps around the hand area of the left primary motor cortex (M1) and assessing the presence/absence of motor-evoked potentials (MEP amplitude > 50 µV; belly-tendon montage). On average, rMT (mean ± SD) was: 45±9% for the vPAS-75 session, 45.2±9.9% for the vPAS-100 session, and 45±9.2% for the vPAS-125 session (no differences among sessions: *F*_2,48=_0.21, *p*=.812). For M1 stimulation, the coil was held tangentially to the scalp and tilted 45° relative to the midline, inducing posterior-to-anterior current flow (Orth & Rothwell, 2004). Electromyography (EMG) was recorded using a Digitimer D360 amplifier (Digitimer Ltd, Welwyn Garden City, UK) connected to a CED micro1401 A/D converter (Cambridge Electronic Devices, Cambridge, UK).

For left V1 stimulation, the coil was placed 2 cm anterior and 0.5 cm lateral to the inion, roughly over the O1 scalp site, following established literature to effectively target V1 (Silvanto et al., 2005). The coil was held tangentially to the scalp with the handle pointing horizontally to the right, inducing a lateral-to-medial current flow (Kammer et al., 2001).

The stable TMS coil placement and position during the experimental sessions were continuously monitored using the Navigated Brain Stimulation (NBS) system (Nexstim™, Helsinki, Finland) based on infrared-based frameless stereotaxy.

### 2.4. Visual-evoked potentials (VEP) and electroencephalography (EEG) recording

VEPs were elicited by black-and-white, left- and right-lateralized pattern reversal checkerboards (digital Michelson contrast: 100%, mean luminance: 200 cd/m^2^, 7.4° × 7.4° visual angle; same visual stimuli as in vPAS). Stimuli were presented at a rate of 2.5 Hz (i.e., reversals occurring every 400 ms), alternating between the left and right visual hemifields. At the onset of each checkerboard reversal, a trigger was delivered via a parallel port to the EEG system for VEP recording (**Figure 1b**). Participants completed 800 trials (400 in the left VF and 400 in the right VF), divided into 2 blocks of 400 trials each (200 trials per hemifield per block), with a 30-sec pause between blocks. Each block lasted approximately 3 minutes. Participants were instructed to fixate the centre of the screen, where a red cross appeared throughout the entire task. To obtain a stable pupil diameter, each participant adapted to ambient room lighting for 10 min before VEP recording. Stimulus presentation and EEG trigger delivery were under computer control (E-Prime 3.0, Psychology Software Tools, Inc.). Continuous EEG data were recorded from a 60-channel EEG cap (EasyCap, BrainProducts GmbH, Munich, Germany) connected to a Nexstim EEG system (Helsinki, Finland). Two forehead electrodes served as ground and reference. Two electro-oculographic channels (EOG) were placed over the right eyebrow and the left cheekbone to detect and monitor ocular artifacts associated with eye movements and blinking. Electrode impedance was kept below 5 kω, and data were acquired at 1450 Hz.

### 2.5. VEP pre-processing

EEG data were pre-processed and analysed using Python (v 3.11.11). Continuous recordings were band-pass filtered between 0.1 and 40 Hz. Channels with poor signal quality were identified via visual inspection of time series and power spectral density plots and interpolated using spherical spline interpolation. Independent component analysis (ICA; FastICA algorithm, retaining components explaining 99% of variance) was performed on a 1-Hz high-pass-filtered copy of the data to optimize decomposition stability. We identified components corresponding to ocular, muscular, or other non-neural artifacts by visually inspecting component topographies and time courses, and subtracted them from the original filtered data to obtain artifact-corrected continuous EEG. Cleaned data were re-referenced to the average reference. Epochs were extracted from -100 to 300 ms relative to stimulus onset, and baseline-corrected using the pre-stimulus window (−100 to 0 ms). Epochs with peak-to-peak amplitudes exceeding 100 µV at any channel were rejected, and the remaining epochs were visually inspected for residual artifacts.

After data cleaning, VEPs were computed by averaging artifact-free epochs separately for left and right VF stimulation. Peak amplitudes and latencies were automatically extracted within predefined latency windows centred on the expected N75, P100, and N135 components (55–95, 80–120, and 110–155 ms, respectively; e.g., Di Russo et al., 2005). For each component, values were extracted from the occipital electrode cluster contralateral to the stimulated visual hemifield: a left cluster (P3, P1, PO3, O1) for right visual hemifield presentation and a right cluster (P2, P4, PO4, O2) for left visual hemifield presentation (**Figure 1b**, right panel). Within each latency window, the peak was identified as the most negative deflection for N75 and N135 and the most positive deflection for P100. As a quality check, two experimenters (FC, GG) visually inspected the automatically identified peaks and manually adjusted them when necessary. Manual adjustments were required in fewer than 5% of cases.

### 2.6. Statistical analysis

Each dependent variable (N75, P100, and N135 amplitudes) was analysed separately using repeated-measures ANOVA (rmANOVA) with visual Hemifield (left, right), Time (pre, post), and ISI (75, 100, 125) as within-subject factors. Significant Hemifield x Time x ISI interactions were further explored by conducting separate rmANOVAs for each visual hemifield.

We computed the inter-hemispheric transfer time (IHTT) of the N75 component as the difference between the N75 latency in the ipsilateral and contralateral hemisphere (relative to the visual stimulus hemifield). Left-to-right and right-to-left IHTT were analysed separately via rmANOVAs with Time and ISI as within-subject factors.

Additionally, exploratory generalized linear regression models (Gaussian distribution; identity link function; Gallucci, 2019) were used to examine the association between baseline (pre-vPAS) N75 amplitude (predictor) and the magnitude of vPAS aftereffects (post-minus pre-vPAS difference) on the same component.

Data distribution normality was verified using the Shapiro-Wilk test and Q-Q plots, while sphericity was assessed with Mauchly’s test. When significant, post-hoc comparisons were performed using Tukey’s honest significant difference correction for multiple comparisons. Effect sizes were reported as partial eta-squared (ηp^2^) for rm-ANOVAs and Cohen’s *d* for paired t-tests.

Unless otherwise specified, in the **Results** sections, data are presented as mean ± standard error. Statistical significance was set at *p<.*05. Statistical analyses were performed using the software Jamovi (version 2.7.6) (The Jamovi Project, 2025).

### 2.7. Results

#### 2.7.1. VEP amplitude and IHTT

##### N75

The rmANOVA revealed significant main effects of Hemifield (F_1,24=_11.34, *p*=.003, ηp²=.321), as well as significant Hemifield x Time (F_1,24=_4.37, *p*=.047, ηp²=.154) and Hemifield x Time x ISI (F_2,48=_3.67, *p*=.033, ηp²=.133) interactions. All other main effects and interactions failed to reach statistical significance (all Fs<2.04, all ps>.166; **Supplemental Table 1**).

For N75 elicited by right-sided visual stimuli, no significant main effects or interactions emerged (Time: F_1,24=_0.01, *p*=.909, ηp²=.001, ISI: F_2,48=_0.42, *p*=.657, ηp²=.017, Time x ISI interaction: F_2,48=_0.59, *p*=.556, ηp²=.024, **Figure 2a, left panel**). Converserly, for left-sided visual stimuli, significant effects of Time (F_1,24=_5.61, *p*=.026, ηp²=.190) and Time x ISI interaction were found (F_2,48=_4.50, *p*=.016, ηp²=.158), whereas the main effect of ISI did not reach statistical significance (F_2,48=_0.16, *p*=.852, ηp²=.007). Post-hoc comparisons showed a significant reduction in N75 amplitude selectively following the vPAS-75 (pre: -1.37±0.10 μV *vs.* post: -1.05±0.08 μV; t_24=_-3.51, *p_Tukey_*_=_.02, d=-0.70). No significant changes were observed following vPAS-100 (1.30±0.11 μV *vs.* - 1.19±0.14 μV; t_24=_-1.22, *p_Tukey_*_=_.826, d=-0.24) or vPAS-125 (−1.26±0.10 μV vs. -1.26±0.12 μV; t_24=_- 0.12, *p_Tukey_*_=_.99, d=-0.02; **Figure 2a, right panel**).

**Figure 2.**
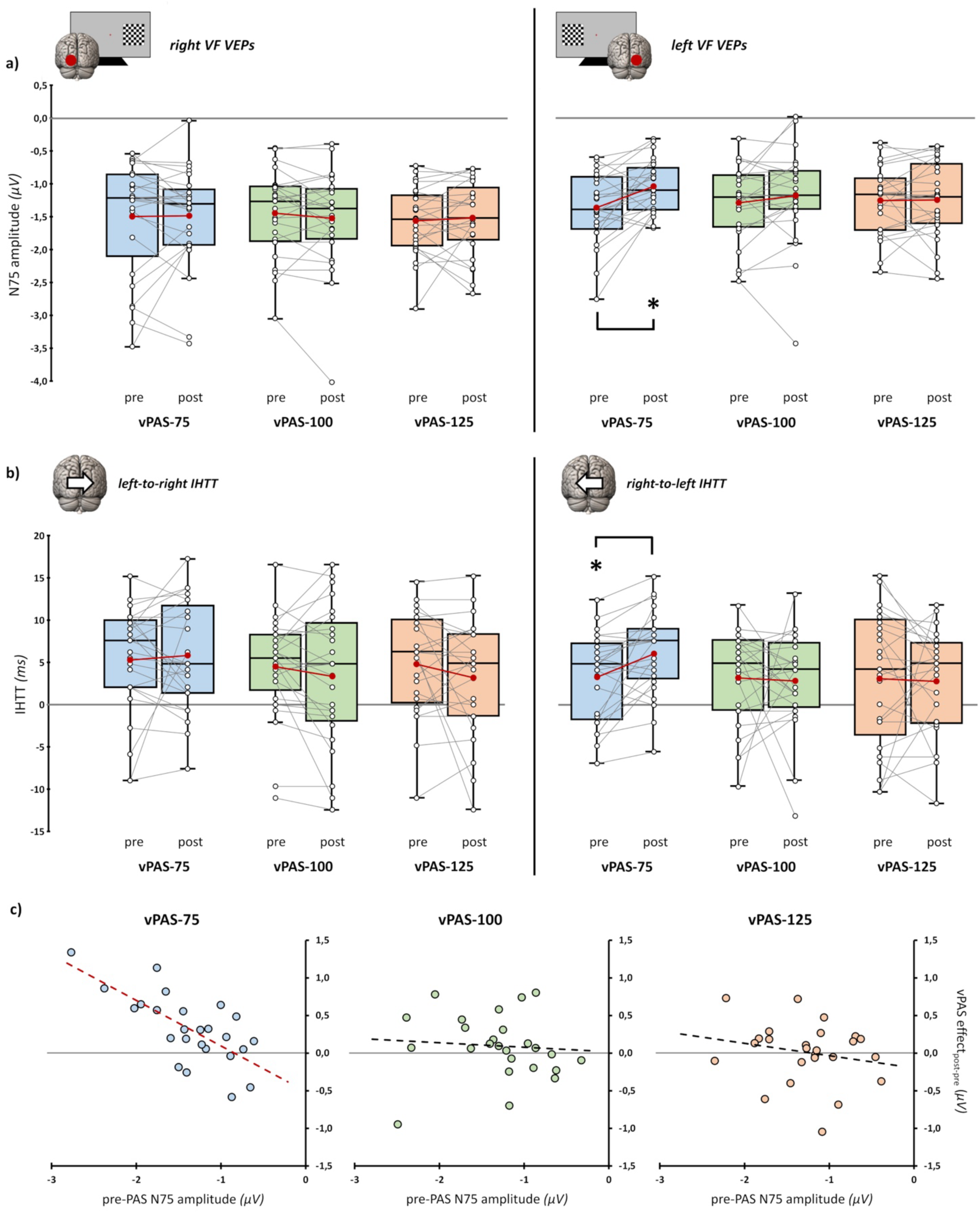
Experiment 1 neurophysiological results. (**a**) N75 amplitude in the three experimental conditions (blue boxplots: vPAS-75; green boxplots: vPAS-100; red boxplots: vPAS-125), before and after PAS administration. In the left panel, amplitude values for stimuli presented in the right visual field (VF); in the right panel, values for stimuli presented in the left VF. (**b**) N75 inter-hemispheric transfer time before and after vPAS (IHTT, left panel: left-to-right IHTT, right panel: right-to-left IHTT). (**c**) Scatterplots between vPAS after-effects (i.e., post-minus pre-vPAS N75 amplitude) and the amplitude of the same component recorded in the respective baseline. Dashed lines indicate the linear regression’s fitted line. In box-and-whisker plots, red dots and lines indicate the means of the distributions. The centre line denotes their median values. Black- and-white dots and grey lines show individual participants’ scores. The box contains the 25th to 75th percentiles of the dataset. Whiskers extend to the largest observation falling within 1.5 × the interquartile range from the first/third quartile. Asterisks indicate statistical significance in post-hoc tests corrected for multiple comparisons (* = pTukey<.05; ** = pTukey<.01; *** = pTukey<.001).

##### P100

Only the Time x Hemifield interaction reached statistically significant, indicating that, following vPAS administration, P100 amplitude differed between stimuli presented in the left and right hemifields regardless of the administered protocol (F_1,24=_9.9, *p*=.004, ηp²=.292). No other main effects or interactions were significant (all Fs<2.84, all ps>.068) (**Supplemental Figure 1a, Supplemental Table 2**).

##### N135

The main factors and their interactions did not reach statistical significance (all Fs<4.23, all ps>.051; **Supplemental Figure 1b, Supplemental Table 3**).

##### N75 IHTT

A significant Time x ISI interaction was found for signal transfer from right V1 to left V1 (F_2,48=_3.75, *p=*.031, ηp²=.135): IHTT significantly increased following vPAS-75 (6.08±0.98 ms), compared to baseline (pre-vPAS: 3.32±1.03 ms; t_24=_3.52; *p_Tukey_*_=_.019, d=0.70). Instead, no significant pre- vs. post-vPAS differences emerged for vPAS-100 (3.32±1.03 ms vs. 2.87 ± 1.14 ms; t_24=_0.23, *p_Tukey_*_=_.99, d=0.05) or vPAS-125 (3.03±1.58 ms vs. 2.72±1.25 ms; t_24=_0.22, *p_Tukey_*_=_.99, d=0.04; **Figure 2b, right panel**). The factor Time (F_1,24=_0.44, *p=*.516, ηp²=.018) and Protocol (F_2,48=_1.77, *p=*.181, ηp²=0.069) did not reach statistical significance.

Regarding signal transfer from the left to right V1, no significant effects were found (Time: F_1,24=_1.77, *p*=.196, ηp²=.069, ISI: F_2,48=_2.59, *p*=.085, ηp²=.097, Time x ISI interaction: F_2,48=_0.89, *p*=.418, ηp²=.036, **Figure 2b, left panel**).

#### 2.7.2. Predictability of vPAS modulations from baseline

Generalized regression models showed that baseline N75 amplitude significantly predicted vPAS-75-induced modulations on the same component (R^2^=0.235, χ^2^_1=_1.59, *p*=.008). Specifically, more negative baseline N75 amplitudes were associated with greater post-vPAS amplitude reductions (B=-0.298, SE=.11, z=-2.66, *p*=.008). This relation was absent for both vPAS-100 (model fit: R^2^<0.01, χ^2^_1<_0.01, *p*=.989) and vPAS-125 (model fit: R^2^=0.11, χ^2^_1=_0.5, *p*=.097; **Figure 2c**).

### 2.10. Interim conclusion

Results from *Experiment 1* demonstrate that only the vPAS-75 protocol induced significant neurophysiological changes, selectively reducing N75 amplitude elicited by visual stimuli presented in the left (ipsilateral/unconditioned) visual hemifield, while simultaneously increasing IHTT from right to left V1. Furthermore, the baseline N75 amplitude predicted the magnitude of vPAS-75-induced aftereffects on the same component.

## 3. EXPERIMENT 2: behavioural effects of vPAS on contrast detection

### 3.1. Participants

Twenty-six healthy volunteers took part in *Experiment 2*. One participant dropped out after the first session for personal reasons, leaving a final analysed sample of 25 participants (11 males; mean age: 23.8±3 years; mean education: 15.8±1.8 years). All participants were recruited using the same criteria as *Experiment 1*.

### 3.2. Experimental procedure

The same within-subjects experimental design as in *Experiment 1* was adopted: participants completed three experimental sessions, during which vPAS-75, vPAS-100, and vPAS-125 were administered (see *Experiment 1* for details) before and after a contrast detection task. Each session lasted about 1.5 hours. On average, participants’ rMT was: 38.6±7.9% for the vPAS-75, 39.3±7.5% for the vPAS-100, and 39±8% for the vPAS-125, with no differences among sessions (*F*_2,48=_0.52, *p*=.597).

### 3.3. Contrast detection task

The contrast detection task required participants to report the orientation (i.e., horizontal or vertical) of Gabor patches presented in either the left or right visual hemifield while maintaining central fixation.

Gabor patches (spatial frequency: 1 cycle/°, mean luminance: 200 cd/m^2^, dimension: 7.4° x 7.4° visual angle, centre at 8° eccentricity; identical position and dimension as the vPAS checkerboards) were presented at 10 contrast levels ranging from high contrast to low contrast (digital Michelson contrasts: 34.6%, 9.3%, 3.9%, 2.8%, 2.3%, 1.6%, 1.2%, 0.8%, 0.5%, and 0.2%).

Each trial began with a red fixation cross presented at the centre of the screen on a grey background for a jittered duration of 800-1500 ms. Subsequently, the Gabor patch appeared for 100 ms, followed by a red question mark lasting 1500 ms that signalled participants to respond. Participants pressed the left mouse button for horizontal Gabor patches and the right mouse button for vertical Gabor patches. Participants were instructed to respond on every trial, guessing when uncertain (i.e., a 2-alternative-forced-choice task). The trial ended with a 300-ms random visual noise mask.

Participants completed 240 trials (120 verticals, 120 horizontals; trial duration: ∼2.5–3 s) split across six blocks of 40 trials, separated by 30-s pauses. Each contrast level and visual hemifield combination was presented 12 times. Participants completed the first three blocks using their right hand and the remaining blocks using their left. Response buttons remained constant across blocks (i.e., left mouse key for horizontal, right key for vertical Gabor patches). The entire task lasted approximately 12 minutes and was implemented in E-Prime 3.0 (Psychology Software Tools, Inc.).

To ensure comprehension of task rules and response mapping, participants completed two 10-trial practice blocks at the start of each experimental session (one per hand). During practice, only high-contrast Gabor patches were presented (levels 6-10; see **Figure 3**). Participants had to achieve accuracy above 75% in both practice blocks before proceeding to the main experiment.

**Figure 3.**
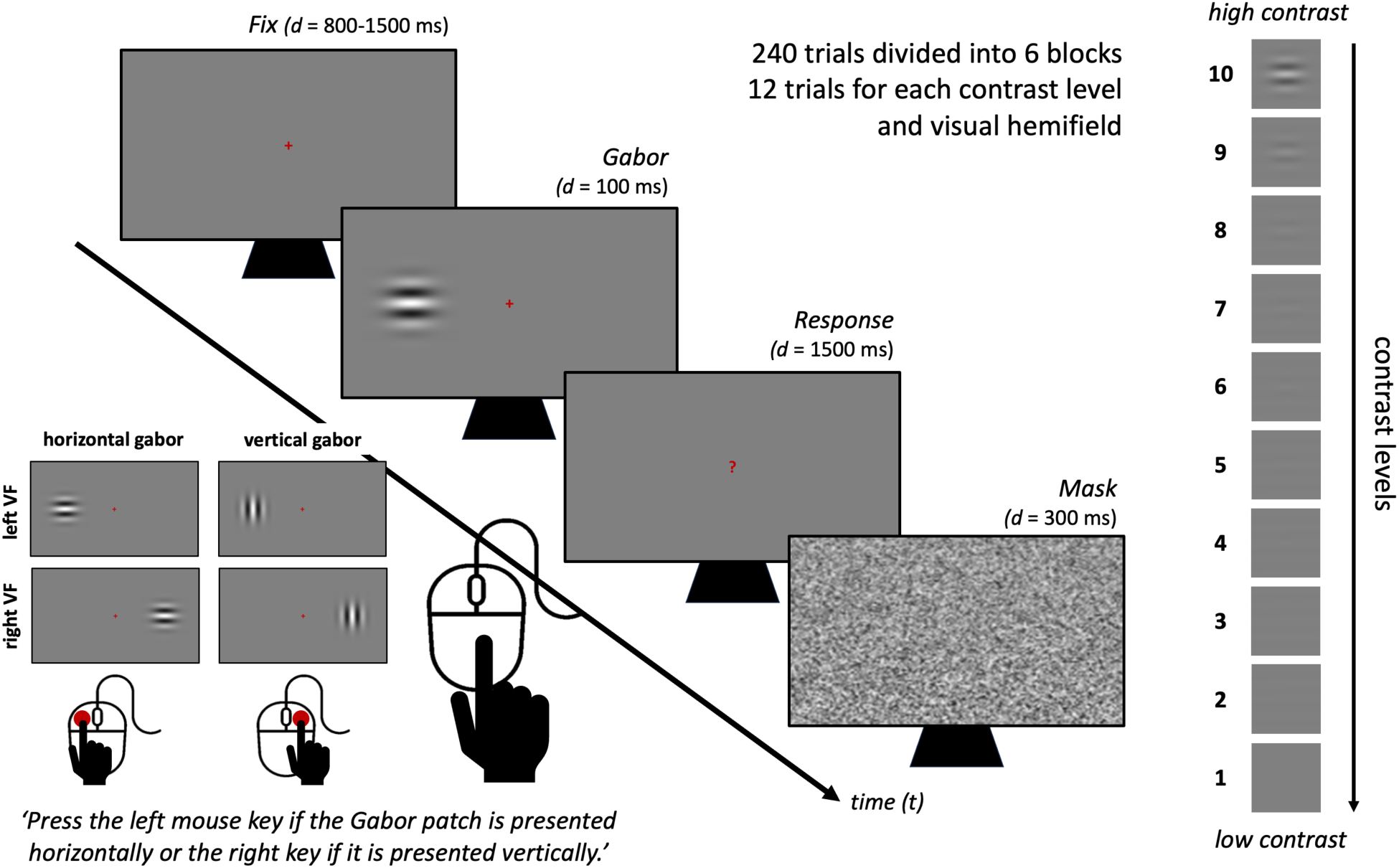
Contrast detection task. Schematic representation of a single trial. On the right, the 10 Gabor patch contrast levels used in the experiment. Depending on the orientation of the Gabor patch (horizontal, vertical) and the visual hemifield of presentation (left vs. right visual field - VF), four different stimulus conditions were possible. Participants responded with their right hand in half of the experimental blocks and with their left hand in the remaining clocks.

Task performance was evaluated using Signal Detection Theory (Green & Swets, 1966) to disentangle perceptual sensitivity (*d-prime – d’*) from response bias (*criterion - c*). We also extracted contrast thresholds and slopes from the fitted logistic function. Psychometric curves were estimated by fitting a generalized linear model (binomial family) in R (R Core Team, 2026) on *dʹ* values normalized to 0–1 (for a similar procedure, see Zazio et al., 2026). We defined the contrast threshold as the contrast level corresponding to the midpoint (50%) of the fitted function, representing the estimated contrast needed for reliable orientation discrimination. We extracted the slope as a secondary index of perceptual precision, with steeper slopes indicating a more abrupt performance gain with increasing contrast. Fitted psychometric curves across conditions are depicted in **Figure 4**.

**Figure 4.**
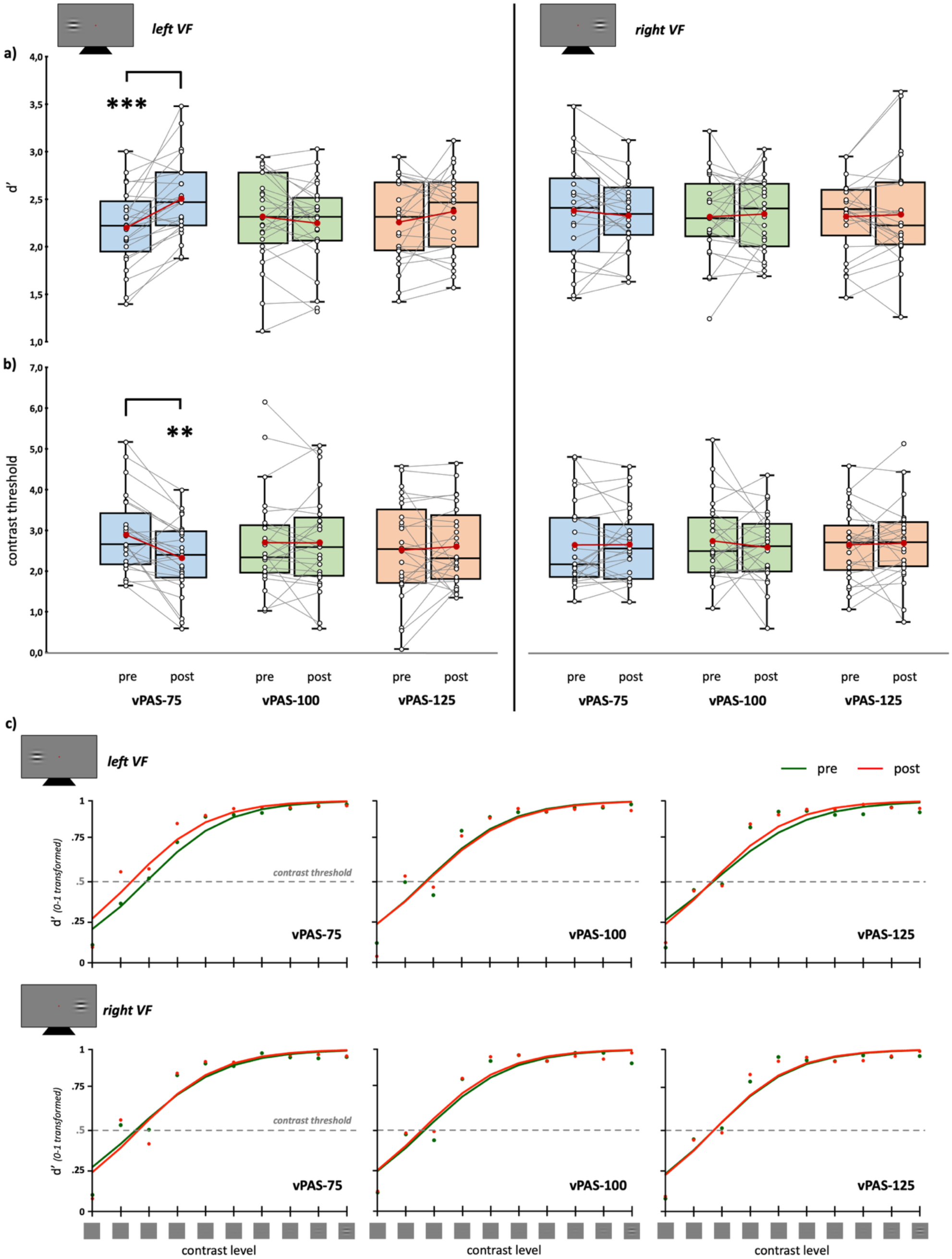
Experiment 2 behavioural results. Perceptual sensitivity *(d’*, **a**) and contrast threshold (**b**) across the three vPAS (blue boxplots: vPAS-75; green boxplots: vPAS-100; red boxplots: vPAS-125), before and after its administration. Left panels display values for stimuli presented in the left visual hemifield (VF), right panels display values for stimuli presented in the right hemifield. Red dots and lines represent group means, where centre lines denote median values. Black-and-white open circles and grey lines show individual participant scores. Box boundaries represent the 25th to 75th percentiles. Whiskers extend to the most extreme data points within 1.5 × interquartile range (IQR) from the first/third quartile. (**c**) Psychometric functions to obtain contrast thresholds, before (green line) and after (red line) each vPAS protocol according to stimulus presentation hemifield (upper panels: left VF, lower panels: right VF). Asterisks indicate statistical significance in post-hoc comparisons for multiple testing (* = pTukey<.05; ** = pTukey<.01; *** = pTukey<.001).

### 3.4. Statistical analysis

We analysed the data using the same statistical approach as in *Experiment 1*. Each dependent variable (*d’*, *c*, contrast threshold, slope of the fitted function) was analysed separately using rmANOVA with visual Hemifield (left, right), Time (pre, post), and ISI (75, 100, 125) as within-subject factors. Significant triple interactions were explored via separate rmANOVAs for each visual hemifield.

### 3.5. Results

#### Sensitivity (d’)

The rmANOVA showed significant Hemifield x Time (F_1,24=_4.33, *p*=.048, ηp²=.153) and Hemifield x Time x ISI (F_2,48=_5.58, *p*=.007, ηp²=.189) interactions. All other main effects and interactions did not reach statistical significance (all Fs<2.42, all ps>.133; **Supplemental Table 4**).

For stimuli presented in the right visual hemifield, no significant effects emerged (Time: F_1,24=_0.15, p=.702, ηp²=.006, ISI: F_2,48=_0.02, p=.984, ηp²=.001, Time x ISI: F_2,48=_0.41, p=.663, ηp²=.017; **Figure 4a, right panel**), whereas for left-sided visual stimuli, significant effects of Time (F_1,24=_5.51, *p*=.027, ηp²=.187) and Time x ISI (F_2,48=_10.38, *p*<.001, ηp²=.302) were found. Post-hoc comparisons showed a significant increase in *d’* selectively after vPAS-75 (2.20±0.08 vs. 2.51±0.08; t24=4.87, *p*_Tukey<_.001, d=0.974). No significant changes were observed following vPAS-100 (2.25±0.09 vs. 2.32±0.09; t24=1.04, *p*_Tukey=_.9, d=0.208) or vPAS-125 (2.26±0.08 vs. 2.37±0.09; t_24=_1.49, *p*_Tukey=_.672, d=0.299; Figure 4a, left panel).

#### Response criterion (c)

No significant main effects or interactions emerged (all Fs<3.67, all ps>.067; **Supplemental Figure 2a, Supplemental Table 5**).

#### Contrast threshold

The rmANOVA revealed a significant Hemifield x Time x ISI interaction (F_2,48=_3.47, *p*=.039, ηp²=.126), whereas no other main effects or interactions reached significance (all Fs<2.88, all ps>.066; **Supplemental Table 6**).

No significant effects emerged for stimuli presented in the right hemifield (Time: F_1,24=_0.15, p=.702, ηp²=.006, Protocol: F_2,48=_0.02, p=.984, ηp²=.001, Time x Protocol: F_2,48=_0.41, p=.663, ηp²=.017; **Figure 4b, right panel**). Converserly, for the left hemifield, a significant Time x ISI interaction was observed (F_2,48=_7.29, *p*=.002, ηp²=.233): post-hoc comparisons showed a significant reduction in contrast threshold selectively following vPAS-75 (2.91±0.19 vs. 2.33±0.18; t_24=_-4.14, *p*_Tukey=_.005, d=-0.827), but not after vPAS-100 (2.71±0.24 vs. 2.71±0.24; t_24=_-0.02, *p*_Tukey=_.99, d=0.005) or vPAS-125 (2.54±0.24 vs. 2.62±0.19; t_24=_0.47, *p*_Tukey=_.99, d=0.095; **Figure 4b, left panel**).

#### Slope

No main effects or interactions reached statistical significance (all Fs<1.71, all ps>.204; Supplemental Figure 2b, Supplemental Table 7).

### 3.6. Interim conclusion

Results from *Experiment 2* confirmed the behavioural efficacy of vPAS-75 (but not vPAS-100 or vPAS-125), demonstrating a selective increase in perceptual sensitivity (d’) and a lower contrast threshold following vPAS-75 administration. Crucially, these behavioural improvements occurred in the absence of changes in perceptual precision (slope) or response criterion (*c*), confirming a genuine gain in visual contrast sensitivity.

## 4. EXPERIMENT 3: Hemispheric vPAS specificity and visuomotor interhemispheric transfer delays

### 4.1. Participants

Twenty-eight healthy volunteers took part in *Experiment 3*. Two participants did not complete all sessions, whereas one participant was excluded due to recording issues, leaving the final analysed sample of 25 participants (10 males; mean age: 21.5±2.9 years; mean education: 13.5±1.9 years). We recruited all participants using the same criteria as in *Experiments 1* and *2*.

### 4.2. Experimental procedure

We used the same within-subjects experimental design as in the previous experiments: participants underwent two experimental sessions in which vPAS targeting either the left or right V1 was administered before and after the contrast detection and Poffenberger tasks. The order of stimulation sessions and behavioural tasks was counterbalanced across participants, with task order held constant within participants across sessions. Each session lasted approximately 1.5 hours.

In *Experiment 3,* only the vPAS-75 was administered. This protocol (hereafter named left-vPAS) was compared with a mirrored version in which both visual stimulus presentation and TMS delivery were reversed (right-vPAS). Specifically, for right-vPAS, the visual stimulus was presented in the left hemifield (8.0° to the left of fixation) and the TMS pulse was delivered over right V1 (approximately over the O2 electrode, identified anatomically following the procedure described in *Experiment 1*). rMT was estimated in every session from the M1 of the hemisphere targeted by vPAS. On average, participants’ rMT was 42.6±6.8% for the left-vPAS session and 45±8.3% for the right-vPAS session, with significantly lower rMT values observed for the dominant hemisphere (t_24=_-2.57, p=.016). All other PAS parameters were identical to those described for *Experiments 1* and *2*.

To reduce the overall duration of the contrast detection task and allow for the administration of the Poffenberger task within the time window of PAS aftereffects (e.g., Guidali et al., 2021b; Suppa et al., 2017), 6 contrast levels were tested in this experiment based on the psychometric results of *Experiment 2* (see **Figure 4c**): 34.6%, 1.6%, 1.2%, 0.8%, 0.5%, and 0.2%. Participants completed 192 trials (96 vertical, 96 horizontal Gabor) across four blocks of 48 trials, separated by 30-sec pauses. Each contrast level and visual hemifield combination was presented 16 times. Participants completed the first two blocks using their right hand and the remaining blocks using their left. The entire task lasted approximately 8 minutes. As for *Experiment 2*, perceptual sensitivity (*d’*), response criterion (*c*), contrast threshold, and psychometric slope were extracted and analysed. Fitted psychometric curves are depicted in **Figure 6**.

**Figure 5.**
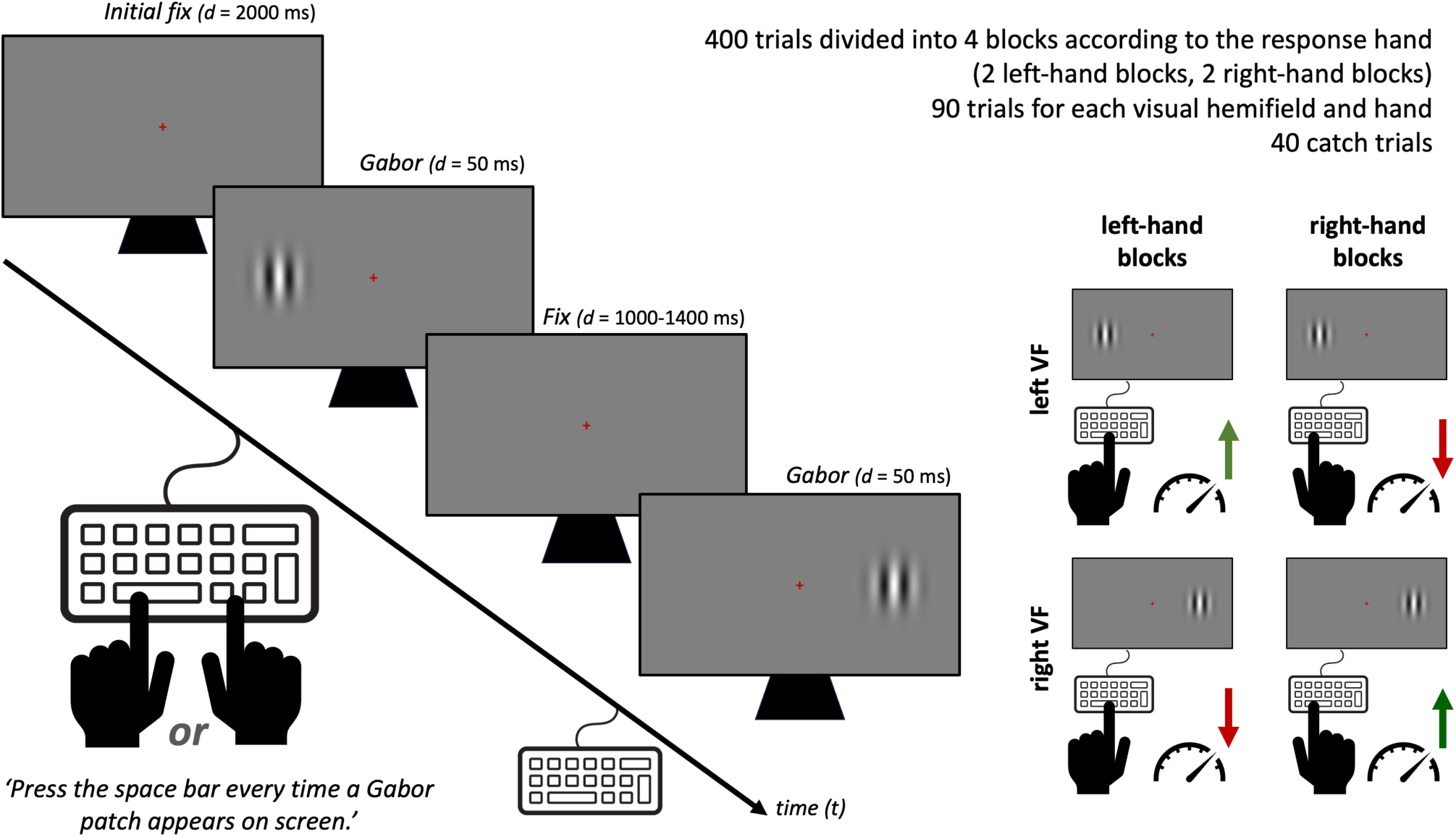
Poffenberger task. Schematic representation of trial sequence and experimental conditions in the Poffenberger task’s trials. Left panel: Timeline of a representative trial. Right panel: Classification of the 4 hand-hemifield conditions. Depending on the responding hand (left vs. right) and the visual hemifield of stimulus presentation (left VF vs. right VF), trials were categorized as uncrossed (green arrows: faster RTs) or crossed (red arrows: slower RTs due to the additional latency required for interhemispheric transfer). Because crossed trials require interhemispheric transfer, they are typically associated with longer reaction times than uncrossed trials.

**Figure 6.**
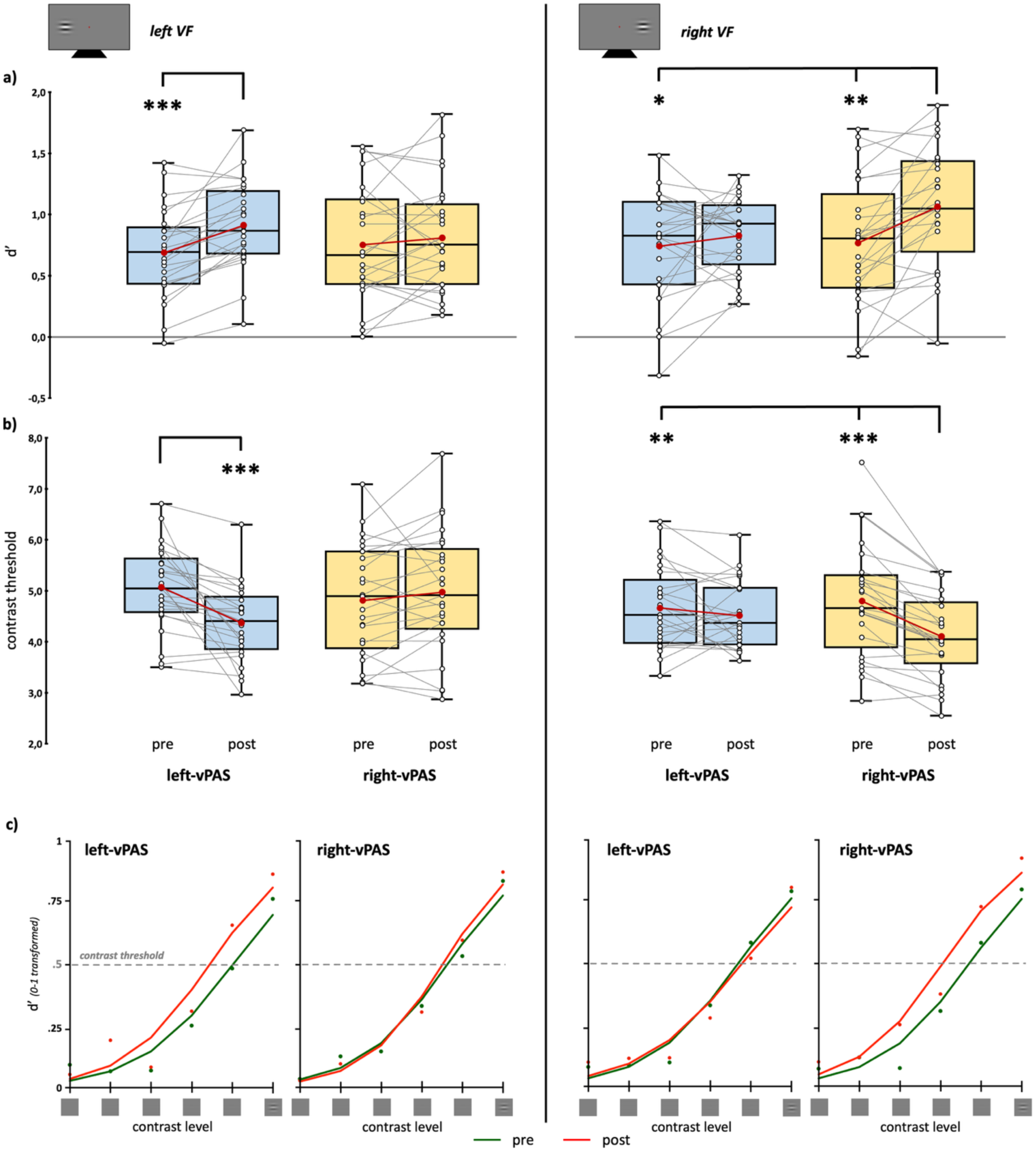
Experiment 3 behavioural results at the contrast detection task. Perceptual sensitivity (d’, **a**) and contrast threshold (**b**) in the two experimental conditions (blue boxplots: left-vPAS; yellow boxplots: right-vPAS), before (pre) and after (post) PAS administration. Left panels display values for stimuli presented in the left visual hemifield (VF); right panels display values for stimuli presented in the right hemifield. Red dots and lines represent means, where centre lines denote median values. Black-and-white open circles and grey lines show individual participant scores. Box boundaries represent the 25th to 75th percentiles. Whiskers extend to the most extreme data points within 1.5 × IQR from the first and third quartiles. (**c**) Psychometric functions from logistic fitting to obtain contrast thresholds, before (green line) and after (red line) each vPAS protocol (left-vPAS and right-vPAS for left visual hemifield - VF; right panels: left-vPAS and right-vPAS for right hemifield). Asterisks indicate statistical significance in post-hoc tests corrected for multiple comparisons (* = pTukey<.05; ** = pTukey<.01; *** = pTukey<.001).

### 4.3. Poffenberger task

Participants were instructed to maintain central fixation and press the space bar as quickly as possible whenever a Gabor patch appeared in the left or right hemifield. Gabor patches had the same spatial frequency, dimensions, and position as those used in the contrast detection task. All stimuli were vertically oriented and presented at a digital Michelson contrast level of 80% to ensure constant supra-threshold visibility.

Each block began with a red fixation cross presented on a grey background at the centre of the screen for 2000 ms. Subsequently, trials began: a Gabor patch appeared for 50 ms, followed by the fixation cross presented for a jittered duration of 1000-1400 ms. As soon as the Gabor appeared, participants responded by pressing the space bar with their index finger (**Figure 5**).

Participants completed 400 trials across four blocks of 100 trials, separated by 30-sec pauses. Participants responded with their right index finger in the first and third blocks, and with their left index finger in the second and fourth blocks, resulting in two blocks per responding hand. Across the task, 180 Gabor patches were presented in each visual hemifield. Overall, for each stimulus location, four hand-hemifield conditions were evaluated: 2 uncrossed conditions (Right Hand/Right VF and Left Hand/Left VF) and 2 crossed conditions (Right Hand/Left VF and Left Hand/Right VF). In addition, 40 catch trials (no stimulus presentation) were included. Average accuracy on catch trials was above 80%. The entire task lasted approximately 8 minutes and was implemented in E-Prime 3.0 (Psychology Software Tools, Inc.).

RTs were recorded for each trial. Trials with anticipatory (< 150 ms) or delayed (> 1200 ms) responses were automatically excluded from further analyses (Jensen, 2006). Participants with 50% trials rejections in any experimental session were excluded from further analyses. Based on this criterion, two participants were removed, leaving the final sample of 23 for the Poffenberger task analyses. Mean RTs were calculated separately for each combination of visual hemifield and responding hand. As the main dependent variable, the crossed-uncrossed difference (CUD) was calculated separately for each visual hemifield as follows (Chaumillon et al., 2014; Marzi et al., 1991):

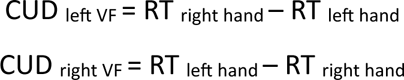

Positive CUD values reflect the additional processing latency associated with visuomotor interhemispheric transfer (Braun, 1992).

Finally, as a measure of overall task performance, total error rates (i.e., omissions and false alarms on catch trials) were calculated.

### 4.4. Statistical analysis

Data were analysed following the same statistical approach as *Experiments 1* and *2*.

For the contrast detection task, each dependent variable was analysed separately using rmANOVA with Visual hemifield (left, right), Time (pre, post), and Protocol (left-vPAS, right-vPAS) as within-subject factors. Significant 3-way interactions were further explored by conducting separate rmANOVAs for each visual hemifield.

For the Poffenberger task, CUD values analysed using the same 2 x 2 x 2 rmANOVA model described above. Total error rates were analysed via rmANOVA with Time and Side-protocol as within-subject factors.

Finally, Spearman correlations were computed between v-PAS-induced changes in contrast detection threshold (*ΔContrast threshold* = Post-PAS *minus* Pre-PAS) and changes in CUD (*ΔCUD* = Post-PAS *minus* Pre-PAS), separately for each hemifield and vPAS protocol.

Data distribution normality, sphericity violations, and post-hoc comparisons were treated as described in previous *Experiments*.

### 4.5. Results

#### 4.5.1 Contrast detection task

##### Sensitivity (*d’*)

The rmANOVA showed significant main effects of Time (F_1,24=_14.15, *p*<.001, ηp²=.371), as well as significant Hemifield x Protocol (F_1,24=_4.32, *p*=.049, ηp²=.152), and Hemifield x Time x Protocol (F_1,24=_15.26, *p*<.001, ηp²=.389) interactions. No other effects reached significance (all Fs<1.52, all ps>.229; Supplemental Table 8).

For stimuli presented in the right hemifield, significant effects of Time (F_1,24=_10.66, *p*=.003, ηp²=.308) and a Time x Protocol (F_1,24=_5.15, *p*=.033, ηp²=.177) interaction were found. Post-hoc comparisons showed a significant increase in *d’* selectively following right-vPAS (0.77±0.10 vs. 1.07±0.10; t_24=_4.10, *p*_Tukey=_.002, d=0.82). No significant changes were observed after left-vPAS (0.74±0.09 vs. 0.83±0.06; t_24=_-1.09, *p*_Tukey=_.702, d=-0.217; **Figure 6a, right panel**). For stimuli presented in the left hemifield, significant main effects of Time (F_1,24=_10.10, *p*=.004, ηp²=.296) and a Time x Protocol interaction emerged (F_1,24=_5.72, *p*=.025, ηp²=.192). Post-hoc comparisons showed a significant increase in *d’* only after left-vPAS (0.68±0.07 vs. 0.91±0.07; t_24=_4.81, *p*_Tukey<_.001, d=0.963; right-vPAS = 0.75±0.09 vs. 0.81±0.09; t_24=_0.85, *p*_Tukey=_.83, d=0.17; **Figure 6a, left panel**).

##### Response criterion (*c*)

Significant main effects of Hemifield (F_1,24=_18.67, *p*<.001, ηp²=.437) and Time (F_1,24=_7.10, *p*=.014, ηp²=.228) were observed, with more negative values for stimuli presented in the left compared to the right hemifield, and more positive values before than after vPAS. No other main effects or interactions reached significance (all Fs<2.33, all ps>.14; **Supplemental Figure 3a, Supplemental Table 9**).

##### Contrast threshold

The rmANOVA showed significant main effects of Time (F_1,24=_27.21, *p*<.001, ηp²=.531) and Hemifield (F_1,24=_7.03, *p*=.014, ηp²=.227), along with a significant Hemifield x Time x Protocol (F_1,24=_41.61, *p*<.001, ηp²=.634) interaction. No other effects reached significance (all Fs<3.82, all ps>.062; **Supplemental Table 10**).

For stimuli presented in the right hemifield, significant effects of Time (F_1,24=_19.45, *p*<.001, ηp²=.448) and Time x Protocol (F_1,24=_12.73, *p*=.002, ηp²=.347) were found. Post-hoc comparisons showed a significant reduction in contrast threshold selectively after right-vPAS (4.79±0.22 vs. 4.10±0.16; t_24=_-6.40, *p*_Tukey<_.001, d=-1.279), but not after left-vPAS (4.67±0.16 vs. 4.52±0.13; t_24=_1.09, *p*_Tukey=_.702, d=0.217; **Figure 6b, right panel**).

Converserly, for stimuli presented in the left hemifield, significant effects of Time (F_1,24=_8.97, *p*=.006, ηp²=.272) and Time x Protocol (F_1,24=_13.04, *p*=.001, ηp²=.352) were found. Post-hoc comparisons showed a significant reduction in contrast threshold after left-vPAS (5.07±0.16 vs. 4.37±0.15; t_24=_-4.38, *p*_Tukey<_.001, d=0.875), but not after right-vPAS (4.81±0.22 vs. 4.97±0.24; t_24=_1.17, *p*_Tukey=_.650, d=0.234; **Figure 6b, left panel**).

##### Slope

No main effects or interactions reached statistical significance (all Fs<1.51, all ps>.234; Supplemental Figure 3b, Supplemental Table 11).

#### 4.5.2. Poffenberger task

##### CUD

The rmANOVA showed significant main effects of Time (F_1,22=_8.22, *p*=.009, ηp²=.272) and Hemifield (F_1,22=_4.57, *p*=.044, ηp²=.172), along with a significant Hemifield x Time x Side-protocol interaction (F_1,22=_4.72, *p*=.041, ηp²=.177). All other main effects and interactions failed to reach statistical significance (all Fs<2.09, all ps>.162; **Supplemental Table 12**).

For stimuli presented in the right hemifield, no significant effects emerged (Time: F_1,22=_0.66, p=.426, ηp²=.029, Protocol: F_1,22=_0.21, p=.653, ηp²=.009, Time x Protocol: F_1,22=_0.7, p=.412, ηp²=.031, **Figure 7a, right panel**). Conversely, for stimuli presented in the left hemifield, significant effects of Time (F_1,22=_6.01, *p*=.023, ηp²=.215) and Time x Protocol (F_1,22=_5.95, *p*=.023, ηp²=.213) were found. Post-hoc comparisons showed a significant reduction in CUD selectively following left-vPAS (6.34±2.22 vs. -2.96±2.01; t_22=_-3.63, *p*_Tukey=_.008, d=-.757). Post-left-vPAS CUD values were also significantly lower compared to post-right-vPAS values (4.54±1.78; t_22=_-3.00, *p*_Tukey=_.031, d=-.630; **Figure 7a, left panel**). **<u>Errors</u>.** The rmANOVA showed a significant main effect of Time (F_1,22=_15.32, *p*<.001, ηp²=.410) and a significant Time x Protocol interaction (F_1,22=_5.83, *p*=.025, ηp²=.210), whereas the main effect of Protocol did not reach significance (F_1,22=_4.16, *p*=.054, ηp²=.159). Post-hoc comparisons revealed a significant increase in errors after left-vPAS (23.9±2.63) compared with both the pre-vPAS assessments (left-vPAS: 15.7±2.05; t_22=_3.75, *p*_Tukey=_.006, d=.782; right-vPAS: 17.17±2.45; t_22=_3.86, *p*_Tukey=_.004, d=.805), The difference between post-left-vPAS and post-right-vPAS showed a trend toward statistical significance (16.83±2.29; t_22=_2.77; *p_Tukey_*_=_.051; d=0.577; Figure 7b).

**Figure 7.**
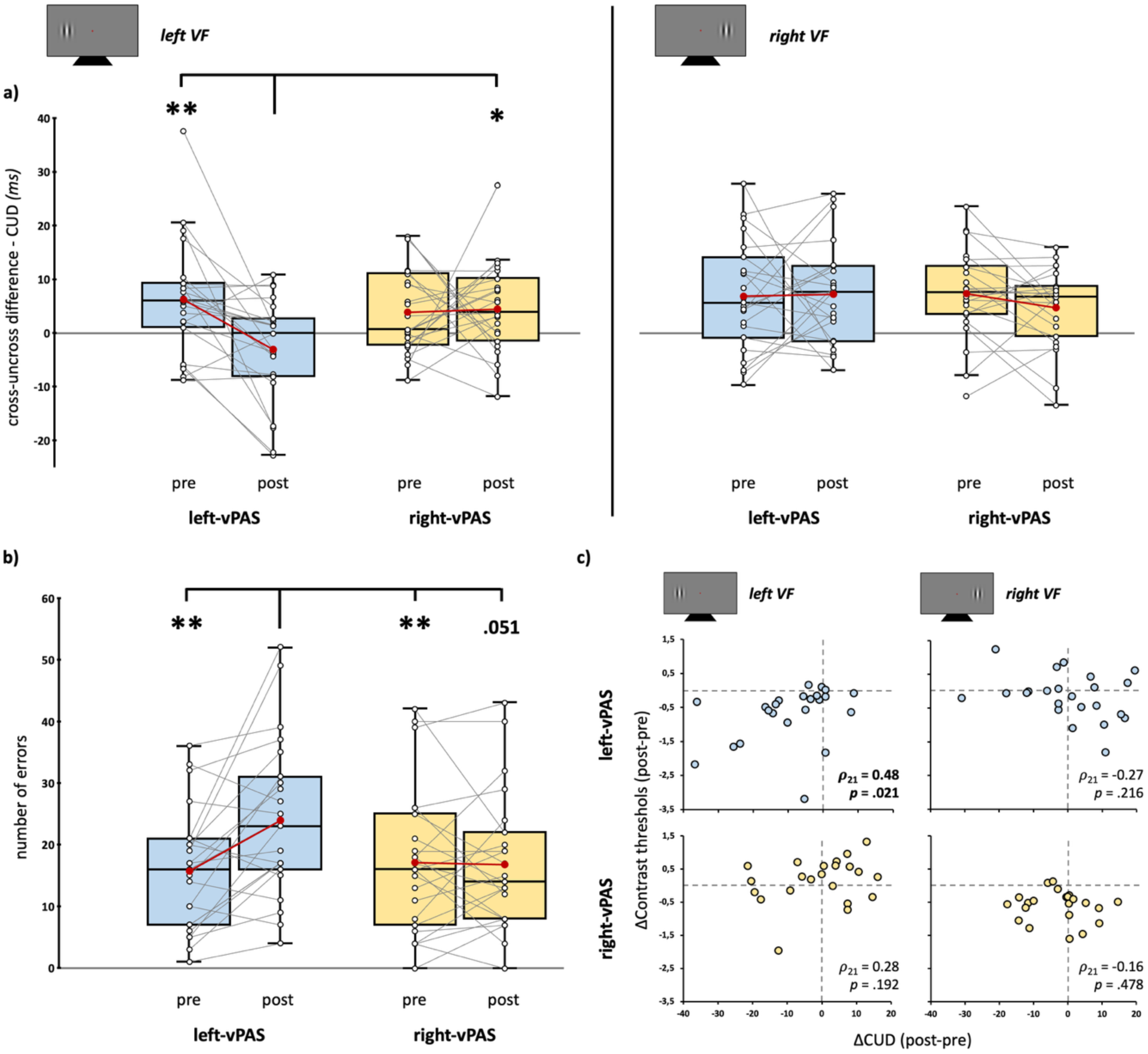
Experiment 3: Poffenberger task and cross-task correlation results. (**a**) Crossed-uncrossed difference (CUD, ms) across experimental conditions (blue boxplots: left-vPAS; yellow boxplots: right-vPAS), before (pre) and after (post) vPAS administration. The left panel displays CUD values for stimuli presented in the left visual hemifield (VF); the right panel displays values for stimuli presented in the right hemifield. (**b**) Number of errors before and after each vPAS protocol. In boxplots, red dots and solid lines represent means, whereas horizontal center lines denote median values. Open circles and grey lines show individual participant scores. Box boundaries represent the 25th and 75th percentiles. Whiskers extend to the most extreme data points within 1.5 x IQR from the first and third quartiles. (**c**) Scatterplots illustrating correlation between vPAS-induced changes in contrast detection task *(ΔContrast threshold*, post–pre) and changes in CUD (*ΔCUD,* post-pre), plotted by visual hemifield (left vs. right VF) and vPAS protocol (left- vs. right-vPAS). Asterisks denote statistical significance in post-hoc test comparisons (* = pTukey<.05; ** = pTukey<.01; *** = pTukey<.001).

#### 4.5.3. Correlations between PAS-induced modulations across behavioural tasks

A significant positive correlation emerged between *ΔContrast threshold* and *ΔCUD* for stimuli presented in the left hemifield following left-vPAS (ρ_21=_.48, *p*=.021), indicating that participants showing a larger PAS-induced reduction in contrast threshold also exhibited greater reduction in CUD. No significant correlations were observed in the right hemifield following left-vPAS (ρ_21=_−.27, *p*=.216), nor for either visual hemifield following right-vPAS (left hemifield: ρ_21=_.28, *p*=.192; right hemifield: ρ_21=_−.16, *p*=.478; **Figure 7c**).

### 4.6. Interim conclusion

Results from *Experiment 3* demonstrate that both left- and right-vPAS-75 protocols induced an hemifield-specific modulation of perceptual sensitivity (*d’*) and contrast thresholds, selectively improving processing of stimuli presented in the visual hemifield contralateral to the stimulated V1. In the Poffenberger task, left-vPAS selectively reduced the CUD for left-sided stimuli, accompanied by an increase in response errors. Crucially, the positive correlation between vPAS-induced changes in contrast threshold and CUD following left-vPAS suggests that the perceptual gain and visuomotor interhemispheric acceleration covaried across participants.

## 5. DISCUSSION

The present study investigated the neurophysiological and functional efficacy of a novel lateralized vPAS protocol targeting human V1. Across three experiments, our results demonstrate that: i) vPAS aftereffects are strictly timing-dependent, occurring selectively at an ISI of 75 ms; ii) these effects induce modulations expressed in electrophysiological and behavioural enhancements in the non-conditioned visual hemifield, ipsilateral to the targeted V1; iii) vPAS-75 modulates both local visual processing and visuo-motor interhemispheric information transfer; iv) the functional pattern of plastic effects varies according to the stimulated hemisphere.

### 5.1. Timing-dependency of vPAS effects

In PAS protocols, the precise temporal relationship governing the two paired inputs represents the key determinant of the magnitude and direction of plasticity-dependent aftereffects (LTP- or LTP-like effects; Guidali et al., 2021b; Suppa et al., 2017). Our findings demonstrate that vPAS was effective exclusively with an ISI of 75 ms between the visual stimulus onset and the TMS pulse (vPAS-75). This strict temporal specificity of vPAS effects may reflect the required coincidence between incoming peripheral visual inputs and TMS-induced cortical activation within V1. Given that the VEP N75 component reflects early activity predominantly generated within the striate cortex (Di Russo et al., 2001), delivering the TMS pulse at a 75-ms latency enables precise temporal overlap between visually evoked and TMS-evoked neural activity in early visual areas, thereby promoting Hebbian associative plasticity. Consistent with this interpretation, vPAS-induced aftereffects were restricted to N75 amplitude and contrast thresholds, two physiological and behavioural markers closely tied to initial V1 processing (Boynton et al., 1999; Di Russo et al., 2001; Shigeto et al., 1998). In contrast, the 100- and 125-ms ISIs align with later stages of visual feedforward processing, during which cortical activity shifts towards extrastriate regions (Di Russo et al., 2001; Pratt, 2011; Šuštar Habjan et al., 2025). In line with Hebbian principles of associative plasticity (Caporale & Dan, 2008) and the timing dependency of PAS effects (Suppa et al., 2017), longer ISIs (>75 ms) reduce the temporal overlap between exogenous TMS activation and endogenous, visually evoked response in V1, rendering associative pairing ineffective.

These results are partially consistent with previous vPAS studies, in which an ISI corresponding to individual P1 latency *minus* 25 ms – yielding an interval close to our fixed 75-ms ISI–successfully modulated VEP habituation, though not VEP amplitude (Ranieri et al., 2019). Notably, Ranieri and coworkers (2019) observed VEP amplitude modulation at an ISI equal to P1 latency *plus* 25 ms, yielding a timeframe comparable to our ineffective vPAS-125 condition. This discrepancy may likely stem from key methodological differences: prior studies administered full-field or central visual stimulation paired with midline TMS over the electrode Oz for bilateral occipital recruitment (Abagnale et al., 2021; Ranieri et al., 2019), whereas our protocol employed lateralized visual stimulation paired with TMS over the contralateral V1. Such spatial configurations distinctly alter the timing and neural dynamics of V1 recruitment, accounting for the unique temporal profile of vPAS aftereffects observed here.

### 5.2. Contralateral effects of lateralized vPAS

The most striking finding is that all electrophysiological and behavioural effects induced by vPAS-75 were selective for stimuli presented in the opposite, unconditioned visual hemifield (ipsilateral to the TMS side), while no modulation emerged for the conditioned visual hemifield targeted by the paired stimulation.

The finding that vPAS can modulate visual processing in the non-stimulated hemisphere suggests that stimulation effects are not confined to the targeted visual cortex and likely involve interhemispheric communication. This evidence aligns with the well-established role of the corpus callosum, particularly its posterior portion (the splenium), in mediating communication between the two visual cortices and supporting the integration of visual information across the visual hemifields (De Lacoste et al., 1985; Pietrasanta et al., 2012). Evidence from patients with unilateral or bilateral V1 lesions also supports this, as interhemispheric transfer of sensory information has been proposed to contribute to the recovery of visual responses after brain damage (Caleo, 2018). Consistent with this role of interhemispheric connections, studies in the motor system have shown that PAS can influence the excitability of contralateral homologous areas (Dickins et al., 2017; e.g., Rizzo et al., 2009; Shin & Sohn, 2011). Similar cross-hemispheric effects have been observed in the visual system using conventional repetitive TMS protocols. For example, Bocci et al. (2011, 2018) showed that unilateral low-frequency rTMS over one visual cortex not only reduced VEP responses to stimuli presented in the contralateral visual hemisphere, but also increased VEP amplitudes for stimuli presented in the ipsilateral hemifield (Bocci et al., 2011, 2018). Although these findings stemmed from a different neurostimulation protocol, they provide clear evidence that modulating one visual cortex can trans-synaptically influence visual responses generated by the contralateral hemisphere. This interpretation is also consistent with TMS-EEG studies showing that unilateral stimulation of the visual cortex can engage distributed cortical networks, with TMS-induced activity extending beyond the stimulated site to regions of the contralateral hemisphere (Garcia et al., 2011).

An alternative, somewhat complementary, hypothesis is that the absence of effects in the visual cortex targeted by the vPAS could reflect a ceiling effect. In healthy individuals, the cortex directly engaged by vPAS may already operate near an optimal level of functional efficiency, limiting how much its activity can be further enhanced (Antal et al., 2006). Under these conditions, associative plasticity may instead manifest more readily in the contralateral visual cortex, potentially through interhemispheric connections.

Regardless of the specific neurophysiological substrate underlying these cross-hemispheric effects, the observed changes included reduced N75 amplitude (*Experiment 1*) and enhanced contrast detection (*Experiment 2*). Both N75 amplitude and visual contrast sensitivity had been associated with activity within V1 (Boynton et al., 1999; Di Russo et al., 2001; Shigeto et al., 1998). The apparent discrepancy between the reduction in VEP amplitude and the improvement in contrast threshold should be interpreted with caution, as several factors may contribute to this dissociation. First, VEPs were elicited using high-contrast pattern-reversal checkerboards (following ISCEV guidelines, Odom et al., 2016), whereas the behavioural task used a range of stimulus contrasts to estimate individual contrast thresholds. This methodological difference may have contributed to the distinct sensitivity of the two measures to vPAS. High-contrast stimuli are optimal for eliciting robust VEP responses, but may provide a restricted dynamic range for detecting plasticity-induced enhancements. Conversely, the lower-contrast stimuli used in the psychophysical task may have increased the sensitivity of visual processing to neuromodulation. Consistent with this possibility, Accornero et al. (2007) showed that the effects of occipital transcranial direct current stimulation on VEPs vary significantly as a function of stimulus contrast (Accornero et al., 2007). Second, reduced VEP amplitude is not necessarily incompatible with improved perceptual performance. For instance, a reduction in C1 amplitude –an early event-related potential component thought to reflect V1 activity (Di Russo et al., 2001; Jeffreys & Axford, 1972) – has been associated with greater behavioural improvements in visual learning paradigms (Ahmadi et al., 2017; Pourtois et al., 2008). Hence, rather than indicating a simple reduction in cortical activity, the N75 pattern found after vPAS may be consistent with more efficient or selective visual processing. Because VEP amplitude primarily reflects the magnitude and synchrony of neural activity across a large population of neurons rather than perceptual efficiency, a reduction in VEP amplitude does not necessarily imply impaired visual processing and may coexist with improved perceptual processing, as shown by enhanced contrast detection after protocol administration.

Taken together, the neurophysiological and behavioural findings suggest that vPAS-75 can modify how early visual cortical responses are recruited during perceptual processing, potentially resulting in a more selective or efficient mode of stimulus coding.

### 5.3. Interhemispheric dynamics and functional hemispheric asymmetry

Beyond these local effects, vPAS-75 also modulated interhemispheric functional interactions. Indeed, the protocol slowed interhemispheric transmission, as indexed by prolonged IHTT for transferring visual information from the right to the left V1 (*Experiment 1*). A related pattern emerged in the Poffenberger task: the typical advantage for uncrossed over crossed responses to stimuli presented in the left, unconditioned visual hemifield was no longer observed after vPAS administration (*Experiment 3)*. Importantly, this attenuation of the crossed-uncrossed difference (CUD) was accompanied by increased response errors, suggesting it did not reflect a genuine improvement in interhemispheric transfer efficiency. Rather, it may indicate an altered balance between hemispheric specialization and interhemispheric integration. Although IHTT and CUD do not measure identical physiological processes, they both provide indirect measures of interhemispheric transfer. IHTT is derived from the summation of neural potentials across neuronal populations (i.e., VEPs) and can therefore be assessed independently of a motor response, providing a direct electrophysiological index of the time required for visual information to traverse the corpus callosum. In contrast, CUD is derived from differences in reaction times between responses to stimuli presented in the two visual hemifields and therefore reflects the consequences of interhemispheric transfer on visuomotor execution. Despite these differences, both measures have been widely employed to assess interhemispheric transfer, capturing this process at different stages of information processing (i.e., early cortical transmission versus visuo-motor integration; Brown et al., 1994).

Taken together, these findings suggest that vPAS exerts a dual influence on both stimulus-specific processing within the non-stimulated visual cortex and the transfer of visual information between the two hemispheres. The improvement in contrast detection, together with the evidence for altered interhemispheric transfer, indicates that vPAS affects local visual processing and interhemispheric communication through partially dissociable mechanisms. One possible interpretation is that the enhanced perceptual performance reflects more efficient processing within the non-stimulated visual cortex, while the concomitant changes in interhemispheric transfer reflect altered cross-callosal integration of visual information. This interpretation is consistent with previous accounts suggesting that increased hemispheric specialization may occur at the expense of cross-hemispheric connectivity (Stephan et al., 2007; Wang et al., 2014).

Notably, this network-level modulation emerged selectively after the vPAS-75 targeted the left V1. Following a protocol targeting the right V1, contrast detection performance improved similarly for stimuli presented in the right VF (corresponding to the non-stimulated left V1), whereas performance on the Poffenberger task was not significantly modulated. Thus, while local perceptual effects emerge regardless of the stimulated hemisphere, the modulation of interhemispheric transfer was specific to the vPAS targeting the left visual system. This asymmetry may reflect well-established hemispheric differences in visual perception and interhemispheric communication. The two hemispheres contribute to visual processing in a complementary manner, with the right hemisphere being preferentially involved in processing of global aspects of visual stimuli, low spatial frequencies, and coordination of spatial relationships, whereas the left hemisphere is more strongly involved in local processing and high spatial frequencies (Hellige, 1996). Robust evidence also demonstrates hemispheric asymmetries in interhemispheric transfer, typically characterized by faster transfer of visual information from the right to the left hemisphere. Such asymmetry has been attributed to structural differences in callosal connections (Marzi et al., 1991), functional specialization (Nowicka et al., 1996), and a relative abundance of fast-conducting, myelinated axons originating in the right relative to the left hemisphere (Barnett & Corballis, 2005).

### 5.4. Limitations

Some methodological and theoretical limitations of the present study warrant consideration. First, because IHTT was not assessed following vPAS targeting the right V1, we could not directly compare electrophysiological interhemispheric modulations between the two hemispheric protocols. Further investigations are required to determine whether the functional asymmetry observed here reflects intrinsic differences in interhemispheric communication or protocol-specific properties of the two stimulation configurations.

Regarding neuronavigation, all three experiments utilized a standard MRI template rather than individual structural scans, which may have affected spatial individualization. However, this standard approach has been successfully validated in previous work to target early visual areas with sub-centimetre precision (Carducci & Brusco, 2012).

Third, TMS intensity was set at 120% of the rMT rather than based on a functional marker of visual cortical excitability, such as the individual phosphene threshold. However, this choice directly aligns with previous vPAS investigations (Abagnale et al., 2021; Ranieri et al., 2019) and is supported by a recent meta-analysis demonstrating a robust correlation between motor and phosphene thresholds (Phylactou et al., 2024).

Finally, given the anatomical proximity and spatial extent of early visual cortices, the TMS-induced electric field likely extended beyond the targeted V1 to partially engage contiguous extra-striate areas, particularly V2 and V3 (see also **Figure 1**). Therefore, although the stimulation site focused primarily on V1, adjacent visual areas may have contributed to the observed physiological and behavioural effects. This limitation is inherent to the spatial resolution of conventional TMS and should be considered when interpreting the present findings as exclusively mediated by V1.

## 6. CONCLUSION

In conclusion, the present findings provide convergent neurophysiological and behavioural evidence that lateralized vPAS-75 can induce plastic changes in the human visual system, extending beyond the stimulated V1. Crucially, these plastic aftereffects manifest predominantly in the non-stimulated visual cortex (hence concern the visual hemifield ipsilateral to the targeted V1), modulating both local perceptual processing and long-range interhemispheric dynamics. The selective efficacy of the 75-ms ISI highlights the importance of stimulation timing for visual associative plasticity. By demonstrating that non-invasive lateralized stimulation can selectively reshape visual sensitivity and cross-callosal communication, these findings advance our understanding of V1 network dynamics and highlight the translational potential of lateralized vPAS for neurorehabilitation in individuals with unilateral visual field defects resulting from damage to a single cerebral hemisphere.

## CREDIT AUTHOR CONTRIBUTION

**Francesca Crespi:** Conceptualization, Methodology, Software, Investigation, Formal analysis, Data curation, Visualization, Writing – original draft

**Giacomo Guidali:** Conceptualization, Methodology, Software, Investigation, Formal analysis, Data curation, Visualization, Supervision, Writing – original draft

**Eleonora Arrigoni:** Methodology, Software, Formal analysis, Visualization, Writing – review and editing

**Nadia Bolognini:** Conceptualization, Methodology, Funding acquisition, Resources, Supervision, Writing – original draft

## ACKNOWLEDGMENTS

We thank Francesca Ornaghi, Daniela Pepe, Alessio Sala, and Davide Zhao for their valuable help with data collection.

## FUNDING

The work was partially supported by the Italian Ministry of Health (Ricerca Corrente 2026).

## COMPETING INTERESTS

The authors declare no competing interests.

## DATA AVAILABILITY

The scripts, raw data, and analyses of this study will be publicly available on an open access repository following manuscript acceptance.

## USE OF AI IN SCIENTIFIC WRITING STATEMENT

During manuscript preparation, the Authors used the software Grammarly Pro to check for grammatical errors and improve sentence structure. After using this tool, the authors reviewed and edited the content as needed and took full responsibility for the publication’s content.

## Notes

### Competing Interest Statement

The authors have declared no competing interest.

