## Supplementary Materials for "Induction of interhemispheric visual plasticity by paired associative stimulation"

### - SUPPLEMENTAL MATERIALS -

*Francesca Crespi, Giacomo Guidali, Eleonora Arrigoni, Nadia Bolognini*

### SUPPLEMENTAL FIGURES

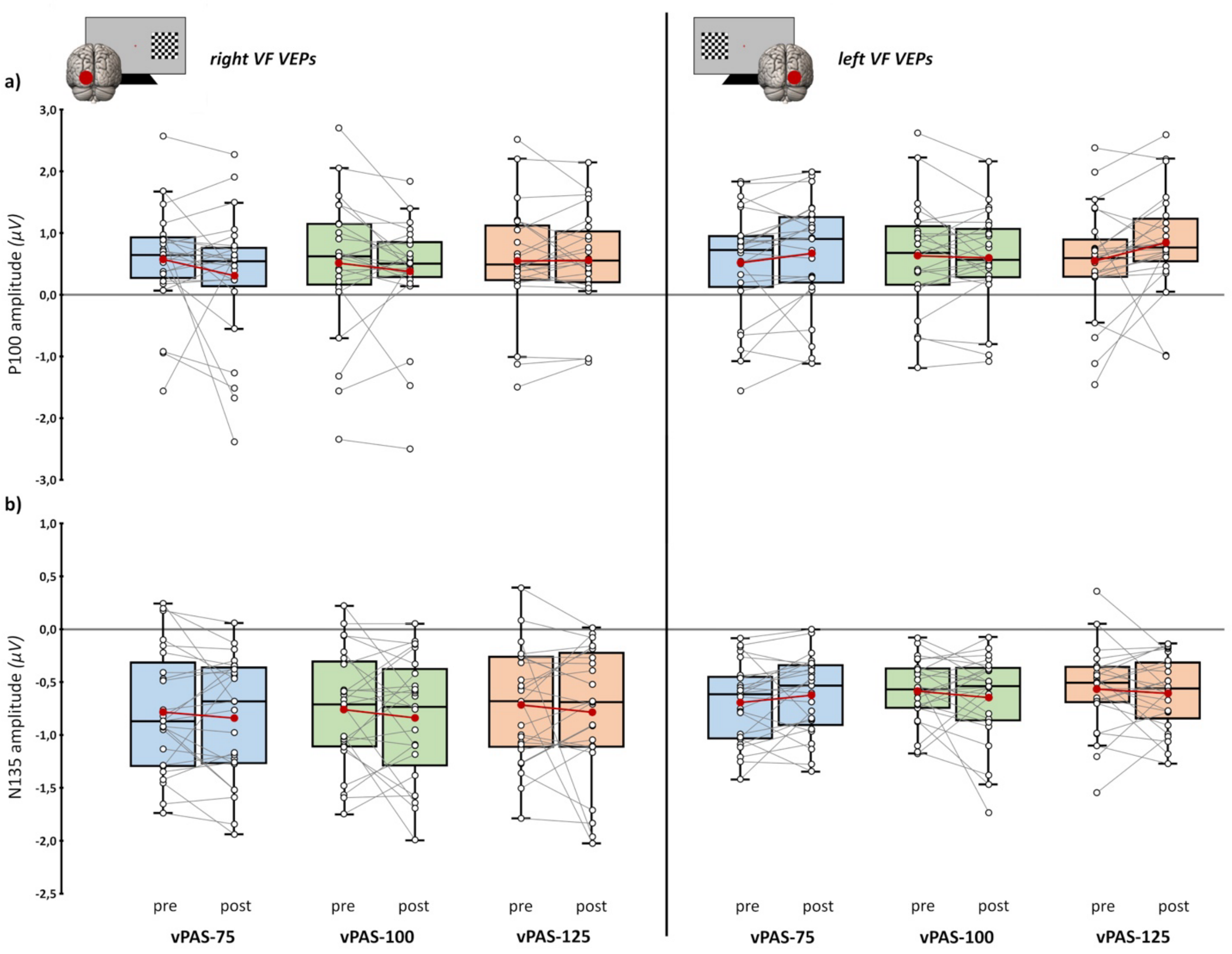


**Supplemental Figure 1.** *Experiment 1 neurophysiological results*. P100 (**a**) and N135 (**b**) amplitude in the three experimental conditions (blue boxplots: vPAS-75; green boxplots: vPAS-100; red boxplots: vPAS-125), before and after PAS administration. In the left panel, amplitude values for stimuli presented in the right visual field (VF); in the right panel, values for stimuli presented in the left VF. In box-and-whisker plots, red dots and lines indicate the means of the distributions. The centre line denotes their median values. Black-and-white dots and grey lines show individual participants' scores. The box contains the 25th to 75th percentiles of the dataset. Whiskers extend to the largest observation falling within 1.5 × the interquartile range from the first/third quartile.

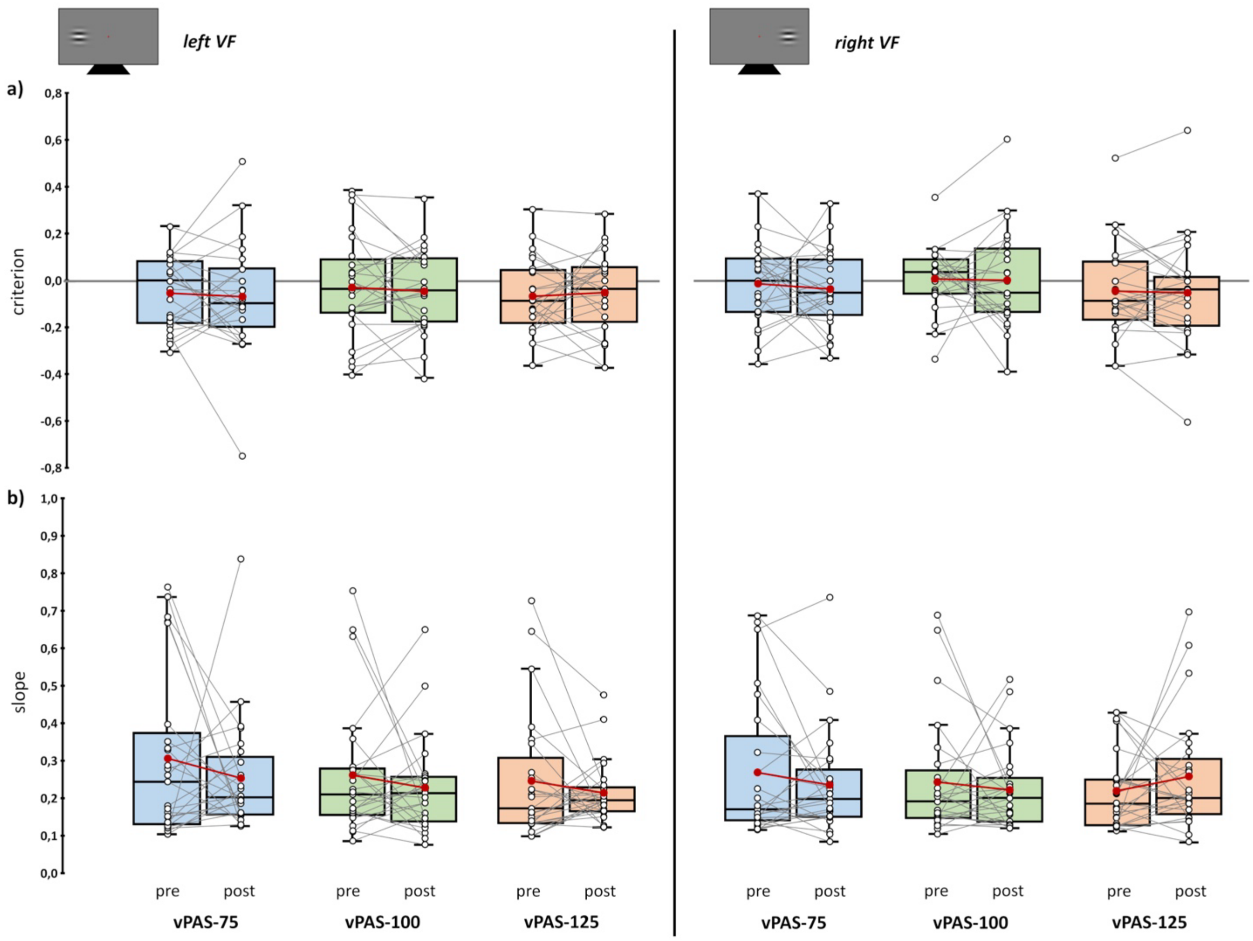


**Supplemental Figure 2.** *Experiment 2 behavioural results*. Response criterion (**a**) and slope of the fitted curve (**b**) in the three experimental conditions (blue boxplots: vPAS-75; green boxplots: vPAS-100; red boxplots: vPAS-125), before and after PAS administration. In the left panel, values for stimuli presented in the left visual field (VF); in the right panel, values for stimuli presented in the right VF. In box-and-whisker plots, red dots and lines indicate the means of the distributions. The centre line denotes their median values. Black-and-white dots and grey lines show individual participants' scores. The box contains the 25th to 75th percentiles of the dataset. Whiskers extend to the largest observation falling within 1.5 × the interquartile range from the first/third quartile.

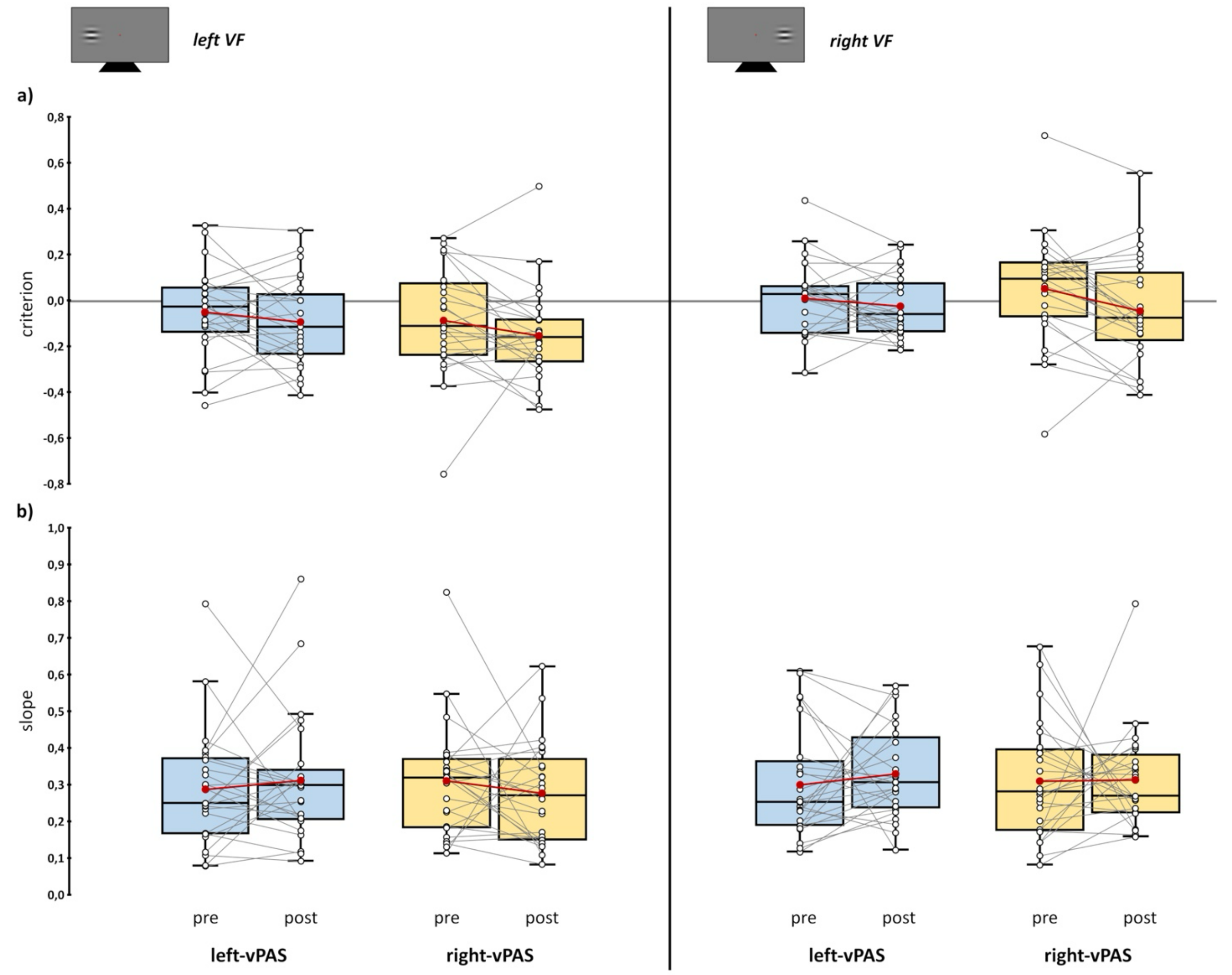


**Supplemental Figure 3.** *Experiment 3 behavioural results*. Response criterion (**a**) and slope of the fitted curve (**b**) in the two experimental conditions (blue boxplots: left-vPAS; yellow boxplots: right-vPAS), before and after PAS administration. In the left panel, values for stimuli presented in the left visual field (VF); in the right panel, values for stimuli presented in the right VF. In box-and-whisker plots, red dots and lines indicate the means of the distributions. The centre line denotes their median values. Black-and-white dots and grey lines show individual participants' scores. The box contains the 25th to 75th percentiles of the dataset. Whiskers extend to the largest observation falling within 1.5 × the interquartile range from the first/third quartile.

### SUPPLEMENTAL TABLES

| Factor/Interaction | F (df1,df2) | *p* | ηp² |
|---|---|---|---|
| ISI | F (2,48) = 0.465 | 0.631 | 0.019 |
| Time | F (1,24) = 2.040 | 0.166 | 0.078 |
| **Hemifield** | **F (1,24) = 11.338** | **0.003** | **0.321** |
| ISI X Time | F (2,48) = 1.724 | 0.189 | 0.067 |
| ISI X Hemifield | F (2,48) = 0.075 | 0.927 | 0.003 |
| **Time X Hemifield** | **F (1,24) = 4.374** | **0.047** | **0.154** |
| **ISI X Time X Hemifield** | **F (2,48) = 3.667** | **0.033** | **0.133** |

**Supplemental Table 1.** vPAS effects in *Experiment 1*: results from the rmANOVA conducted on the N75 amplitude. Statistically significant effects are reported in bold.

| Factor/Interaction | F (df1,df2) | *p* | ηp² |
|---|---|---|---|
| ISI | F (2,48) = 0.371 | 0.692 | 0.015 |
| Time | F (1,24) = 0.079 | 0.781 | 0.003 |
| Hemifield | F (1,24) = 1.228 | 0.279 | 0.049 |
| ISI X Time | F (2,48) = 2.841 | 0.068 | 0.106 |
| ISI X Hemifield | F (2,48) = 0.094 | 0.910 | 0.004 |
| **Time X Hemifield** | **F (1,24) = 9.904** | **0.004** | **0.292** |
| ISI X Time X Hemifield | F (2,48) = 0.963 | 0.389 | 0.039 |

**Supplemental Table 2.** vPAS effects in *Experiment 1*: results from the rmANOVA conducted on the P100 amplitude. Statistically significant effects are reported in bold.

| Factor/Interaction | F (df1,df2) | *p* | ηp² |
|---|---|---|---|
| ISI | F (2,48) = 1.461 | 0.242 | 0.057 |
| Time | F (1,24) = 0.995 | 0.328 | 0.04 |
| Hemifield | F (1,24) = 4.228 | 0.051 | 0.15 |
| ISI X Time | F (2,48) = 0.427 | 0.655 | 0.017 |
| ISI X Hemifield | F (2,48) = 0.158 | 0.854 | 0.007 |
| Time X Hemifield | F (1,24) = 0.843 | 0.368 | 0.034 |
| ISI X Time X Hemifield | F (2,48) = 0.383 | 0.684 | 0.016 |

**Supplemental Table 3.** vPAS effects in *Experiment 1*: results from the rmANOVA conducted on the N135 amplitude.

| Factor/Interaction | F (df1,df2) | *p* | ηp² |
|---|---|---|---|
| ISI | F (2,48) = 0.270 | 0.765 | 0.011 |
| Hemifield | F (1,24) = 0.191 | 0.666 | 0.008 |
| Time | F (1,24) = 2.416 | 0.133 | 0.091 |
| ISI X Hemifield | F (2,48) = 0.198 | 0.821 | 0.008 |

| | | | |
|---|---|---|---|
| ISI X Time | F (2,48) = 2.334 | 0.108 | 0.089 |
| **Hemifield X Time** | **F (1,24) = 4.333** | **0.048** | **0.153** |
| **ISI X Hemifield X Time** | **F (2,48) = 5.579** | **0.007** | **0.189** |

**Supplemental Table 4.** vPAS effects in *Experiment 2*: results from the rmANOVA conducted on the *d'*. Statistically significant effects are reported in bold.

| **Factor/Interaction** | **F (df1,df2)** | **p** | **$\eta p^2$** |
|---|---|---|---|
| ISI | F (2,48) = 2.593 | 0.085 | 0.098 |
| Hemifield | F (1,24) = 3.672 | 0.067 | 0.133 |
| Time | F (1,24) = 0.976 | 0.333 | 0.039 |
| ISI X Hemifield | F (2,48) = 0.531 | 0.592 | 0.022 |
| ISI X Time | F (2,48) = 0.668 | 0.517 | 0.027 |
| Hemifield X Time | F (1,24) = 0.622 | 0.438 | 0.025 |
| ISI X Hemifield X Time | F (2,48) = 0.045 | 0.956 | 0.002 |

**Supplemental Table 5.** vPAS effects in *Experiment 2*: results from the rmANOVA conducted on the response criterion (*c*).

| **Factor/Interaction** | **F (df1,df2)** | ***p*** | **$\eta p^2$** |
|---|---|---|---|
| ISI | F (2,48) = 0.096 | 0.909 | 0.004 |
| Hemifield | F (1,24) = 0.048 | 0.828 | 0.002 |
| Time | F (1,24) = 1.473 | 0.237 | 0.058 |
| ISI X Hemifield | F (2,48) = 0.298 | 0.744 | 0.012 |
| ISI X Time | F (2,48) = 2.881 | 0.066 | 0.107 |
| Hemifield X Time | F (1,24) = 0.799 | 0.380 | 0.032 |
| **ISI X Hemifield X Time** | **F (2,48) = 3.471** | **0.039** | **0.126** |

**Supplemental Table 6.** vPAS effects in *Experiment 2*: results from the rmANOVA conducted on the contrast threshold. Statistically significant effects are reported in bold.

| **Factor/Interaction** | **F (df1,df2)** | ***p*** | **$\eta p^2$** |
|---|---|---|---|
| ISI | F (2,48) = 1.166 | 0.320 | 0.046 |
| Hemifield | F (1,24) = 0.207 | 0.653 | 0.009 |
| Time | F (1,24) = 1.706 | 0.204 | 0.066 |
| ISI X Hemifield | F (2,48) = 0.640 | 0.532 | 0.026 |
| ISI X Time | F (2,48) = 0.747 | 0.479 | 0.03 |
| Hemifield X Time | F (1,24) = 1.183 | 0.288 | 0.047 |
| ISI X Hemifield X Time | F (2,48) = 0.283 | 0.755 | 0.012 |

**Supplemental Table 7.** vPAS effects in *Experiment 2*: results from the rmANOVA conducted on the slope of the fitted function.

| Factor/Interaction | F (df1,df2) | *p* | ηp² |
|---|---|---|---|
| Side | F (1,24) = 0.546 | 0.467 | 0.022 |
| Hemifield | F (1,24) = 1.521 | 0.229 | 0.06 |
| **Time** | **F (1,24) = 14.145** | **<0.001** | **0.371** |
| **Side X Hemifield** | **F (1,24) = 4.317** | **0.049** | **0.152** |
| Side X Time | F (1,24) = 0.106 | 0.748 | 0.004 |
| Hemifield X Time | F (1,24) = 0.863 | 0.362 | 0.035 |
| **Side X Hemifield X Time** | **F (1,24) = 15.258** | **<0.001** | **0.389** |

**Supplemental Table 8.** vPAS effects in the contrast task of *Experiment 3*: results from the rmANOVA conducted on the *d'*. Statistically significant effects are reported in bold.

| Factor/Interaction | F (df1,df2) | *p* | ηp² |
|---|---|---|---|
| Side | F (1,24) = 0.426 | 0.52 | 0.017 |
| **Hemifield** | **F (1,24) = 18.666** | **<0.001** | **0.437** |
| **Time** | **F (1,24) = 7.100** | **0.014** | **0.228** |
| Side X Hemifield | F (1,24) = 2.327 | 0.14 | 0.088 |
| Side X Time | F (1,24) = 1.198 | 0.285 | 0.048 |
| Hemifield X Time | F (1,24) = 0.089 | 0.768 | 0.004 |
| Side X Hemifield X Time | F (1,24) = 0.551 | 0.465 | 0.022 |

**Supplemental Table 9.** vPAS effects in the contrast task of *Experiment 3*: results from the rmANOVA conducted on the response criterion (*c*). Statistically significant effects are reported in bold.

| Factor/Interaction | F (df1,df2) | *p* | ηp² |
|---|---|---|---|
| Side | F (1,24) = 0.010 | 0.92 | <0.001 |
| **Hemifield** | **F (1,24) = 7.031** | **0.014** | **0.227** |
| **Time** | **F (1,24) = 27.208** | **<0.001** | **0.531** |
| Side X Hemifield | F (1,24) = 3.819 | 0.062 | 0.137 |
| Side X Time | F (1,24) = 0.878 | 0.358 | 0.035 |
| Hemifield X Time | F (1,24) = 1.337 | 0.259 | 0.053 |
| **Side X Hemifield X Time** | **F (1,24) = 41.613** | **<0.001** | **0.634** |

**Supplemental Table 10.** vPAS effects vPAS effects in the contrast task of *Experiment 3*: results from the rmANOVA conducted on the contrast threshold. Statistically significant effects are reported in bold.

| Factor/Interaction | F (df1,df2) | *p* | ηp² |
|---|---|---|---|
| Side | F (1,24) = 0.046 | 0.833 | 0.002 |
| Hemifield | F (1,24) = 0.421 | 0.523 | 0.017 |
| Time | F (1,24) = 0.086 | 0.772 | 0.004 |
| Side X Hemifield | F (1,24) = 0.006 | 0.937 | <0.001 |
| Side X Time | F (1,24) = 1.506 | 0.232 | 0.059 |

| | | | |
|---|---|---|---|
| Hemifield X Time | F (1,24) = 0.298 | 0.59 | 0.012 |
| Side X Hemifield X Time | F (1,24) = 0.126 | 0.726 | 0.005 |

**Supplemental Table 11.** vPAS effects in the contrast task of *Experiment 3*: results from the rmANOVA conducted on the slope of the fitted function.

| **Factor/Interaction** | **F (df1,df2)** | ***p*** | **$\eta p^2$** |
|---|---|---|---|
| **Side** | **F (1,22) = 4.572** | **0.044** | **0.172** |
| **Hemifield** | **F (1,22) = 8.223** | **0.009** | **0.272** |
| Time | F (1,22) = 0.553 | 0.465 | 0.025 |
| Side X Hemifield | F (1,22) = 1.520 | 0.231 | 0.065 |
| Side X Time | F (1,22) = 1.087 | 0.308 | 0.047 |
| Hemifield X Time | F (1,22) = 2.092 | 0.162 | 0.087 |
| **Side X Hemifield X Time** | **F (1,22) = 4.726** | **0.041** | **0.177** |

**Supplemental Table 12.** vPAS effects in the Poffenberger task of *Experiment 3*: results from the rmANOVA conducted on the cross-uncross difference (CUD). Statistically significant effects are reported in bold.
